# WNK-SPAK/OSR1 signaling pathway facilitates ictal activity via reduced neuronal chloride extrusion rate

**DOI:** 10.64898/2026.06.10.731153

**Authors:** Volodymyr I. Dzhala, Fu Hung Shiu, Negah Rahmati, Aloe Carroll, Reagan Bae, Kristopher T. Kahle, Kevin J Staley

## Abstract

Seizures upregulate Na^+^-K^+^-2Cl^-^ (NKCC1)-mediated Cl^−^ influx and downregulate K^+^-Cl^-^(KCC2)-mediated Cl^-^ efflux *via* the WNK-SPAK/OSR1 kinases, leading to cytoplasmic chloride ([Cl^-^]_i_) accumulation, reduced GABAergic inhibition and anticonvulsant failure. Early studies found that inhibiting WNK-kinase reduced baseline [Cl^-^]_i_ (E_Cl_) and seizures via increased KCC2 activity. However, increased KCC2 activity alone should not affect E_Cl_ whose determinants are more complex. We determined the net effects of WNK-SPAK/OSR1 pathway inhibitor WNK463 on E_Cl_ and [Cl^-^]_i_ transients during spontaneous ictal-like discharges (ILDs). We found that WNK463 reduced interictal [Cl^-^]_i_ but did not change baseline [Cl^-^]_i_ measured in the presence of TTX. WNK463 enhanced neuronal Cl^-^ extrusion during and after ILDs, before abolishing ILDs. Pharmacological inhibition and targeted siRNA silencing demonstrated that the anti-ictal effects of WNK463 involved both NKCC1 and KCC2. Our data support a mechanism whereby mutual NKCC1 inhibition and KCC2 activation via the WNK-SPAK/OSR1 pathway exert powerful anti-ictal effects by facilitating [Cl^-^]_i_ extrusion during ILDs.

## Introduction

Acute brain injury is often complicated by antiepileptic (AED)-resistant seizures (Liesemer et al., 2011). One potential mechanism of these seizures and AED failure is the inversion of signaling by the inhibitory neurotransmitter GABA in injured neurons. Acute brain injury results in cytotoxic cerebral edema, i.e., an influx of _NaCl_ into neurons and glia (Van Harreveld and Schade, 1959;Steffensen et al., 2015). This _NaCl_ influx results in an elevated baseline [Cl^-^]_i_ and a consequent depolarizing shift in the reversal potential for GABA_A_ receptor-mediated currents (E_GABA_) (van den Pol et al., 1996;Pond et al., 2006;Dzhala et al., 2010;Dzhala et al., 2012;Kahle et al., 2008;Blaesse et al., 2009). Depolarizing GABA responses facilitate neuronal network activity and contribute to initiation of seizures (Dzhala and Staley, 2003;Khazipov et al., 2004). Furthermore high rates of synaptic Cl^-^ influx during seizures stress neuronal Cl^-^ homeostasis (Staley and Proctor, 1999) and shift E_GABA_ to more positive potentials, contributing to prolonged seizures and anticonvulsant-resistance (Khalilov et al., 2003;Dzhala et al., 2010).

After synaptically-mediated Cl^-^ influx, restoration of [Cl^-^]_i_ equilibrium is achieved by the activities of the cation-Cl^-^ co-transporters (CCCs) such as the Na^+^-driven NKCC1, which is biased toward Cl^-^ influx, and the K^+^-driven KCC2, which is biased toward Cl^-^ efflux (Staley, 2024). Depending on the ionic and osmotic gradients across the neuronal cytoplasmic membrane, either antagonizing or stimulating CCCs activity may be a useful therapeutic strategy to reduce [Cl^-^]_i_ accumulation, restore GABAergic inhibition, and suppress seizures (Glykys et al., 2017). Previous studies have demonstrated that inhibition of NKCC1 reduced [Cl^-^]_i_ in injured neurons, enhanced GABAergic inhibition and the efficacy of anticonvulsants (Pond et al., 2006;Dzhala et al., 2008;Dzhala et al., 2012;Dzhala et al., 2024;Nardou et al., 2011;Cleary et al., 2013;Sivakumaran and Maguire, 2016). Conversely, enhancement of KCC2 activity improved [Cl^-^]_i_ extrusion and control of seizures (Gagnon et al., 2013;Moore et al., 2018;Dzhala and Staley, 2021).

Transmembrane Cl^-^ equilibrium is mediated by a Donnan mechanism that involves intra- and extracellular impermeant anions, in which the CCCs comprise the requisite cation-chloride membrane permeability (Glykys et al., 2017;Bahari et al., 2024). Increasing cation-chloride permeability by increasing the maximum velocity of CCCs should not change the baseline [Cl^-^]_i_. However, in injured neurons with elevated [Cl^-^]_i_, increased cation-chloride extrusion might reduce neuronal volume and [Cl^-^]_i_. Further, increasing the maximum velocity of CCCs should increase the ability of neurons to buffer large ictal synaptically-mediated Cl^-^ influxes. The maximum velocity of CCCs is regulated by the WNK-SPAK/OSR1 kinase pathways (Kahle et al., 2006;Melo et al., 2013;Alessi et al., 2014). In hypertonic extracellular conditions, WNK-SPAK/OSR1 signaling phosphorylates NKCC1 (Moriguchi et al., 2005;Vitari et al., 2006), rapidly enhancing its activity and chloride uptake (Côme et al., 2023). In isotonic conditions KCC2 is phosphorylated and inactive, whereas hypotonic conditions promote dephosphorylation and activation of KCC2 (Kahle et al., 2013;Pisella et al., 2019;Watanabe et al., 2019). Therefore, blocking the phosphorylation of KCC2 and NKCC1 via WNK signaling would shift net membrane transport toward maximum Cl^-^ extrusion, preventing excessive [Cl^-^]_i_ accumulation in conditions such as traumatic brain injury and seizures.

The recently identified WNK-kinase inhibitor WNK463 reduces phosphorylation of downstream targets SPAK/OSR1 (Yamada et al., 2017) and thereby inhibits NKCC1 and KCC2 phosphorylation (de Los et al., 2014), resulting in reciprocal inhibition of NKCC1 and stimulation of KCC2 activity. WNK463 reduced KCC2-Thr1007 phosphorylation, hyperpolarized E_GABA_ and limited status epilepticus (Lee et al., 2022). The relative contribution of the WNK-SPAK/OSR1 pathway in the net regulation of the NKCC1 and KCC2 transport activity (velocity) and [Cl^-^]_i_ transients during recurrent seizures was not determined. We determine now the net effects of WNK-SPAK/OSR1 pathway inhibitor WNK463 on [Cl^-^]_i_ elevation and extrusion rates during spontaneous ILDs, and the correlation between ionic and electrographic effects in an *in vitro* model of post-traumatic epilepsy.

## Results

### Chloride transients during recurrent ILDs

Organotypic hippocampal slices from mice expressing the genetically encoded intracellular chloride fluorophore Clomeleon or Super Clomeleon (Grimley et al., 2013) were used as a model of acute traumatic brain injury and epileptogenesis *in vitro* (Dyhrfjeld-Johnsen et al., 2010;Berdichevsky et al., 2012;Dzhala et al., 2012). Simultaneous extracellular field potential recordings and two-photon imaging were performed in the CA1 pyramidal cell layer to monitor neuronal network activity and [Cl^−^]_i_ in individual cells (Figure 1). Spontaneous neuronal network activity was characterized by brief interictal-like discharges (IEDs) and prolonged ictal-like discharges (ILDs) reminiscent of seizure-like activity *in vivo*. In control, during the inter-ictal phase between ILDs, the baseline [Cl^−^]_i_ in individual pyramidal cells varied from 5 to 30 mM and higher (Figure 1a). In line with previous studies (Lillis et al., 2012;Glykys et al., 2014), [Cl^−^]_i_ rapidly increased at the onset of ILDs and during sustained high-frequency ictal-tonic discharges (Figure 1b). Elevated [Cl^−^]_i_ levels often persisted during intermittent ictal-clonic discharges and progressively decreased during secondary after-discharges and termination of ILDs. Postictally, [Cl^−^]_i_ decayed monoexponentially with a decay time constant that varied from 20 to 240 s (Figures 1, 5).

**Figure 1.**
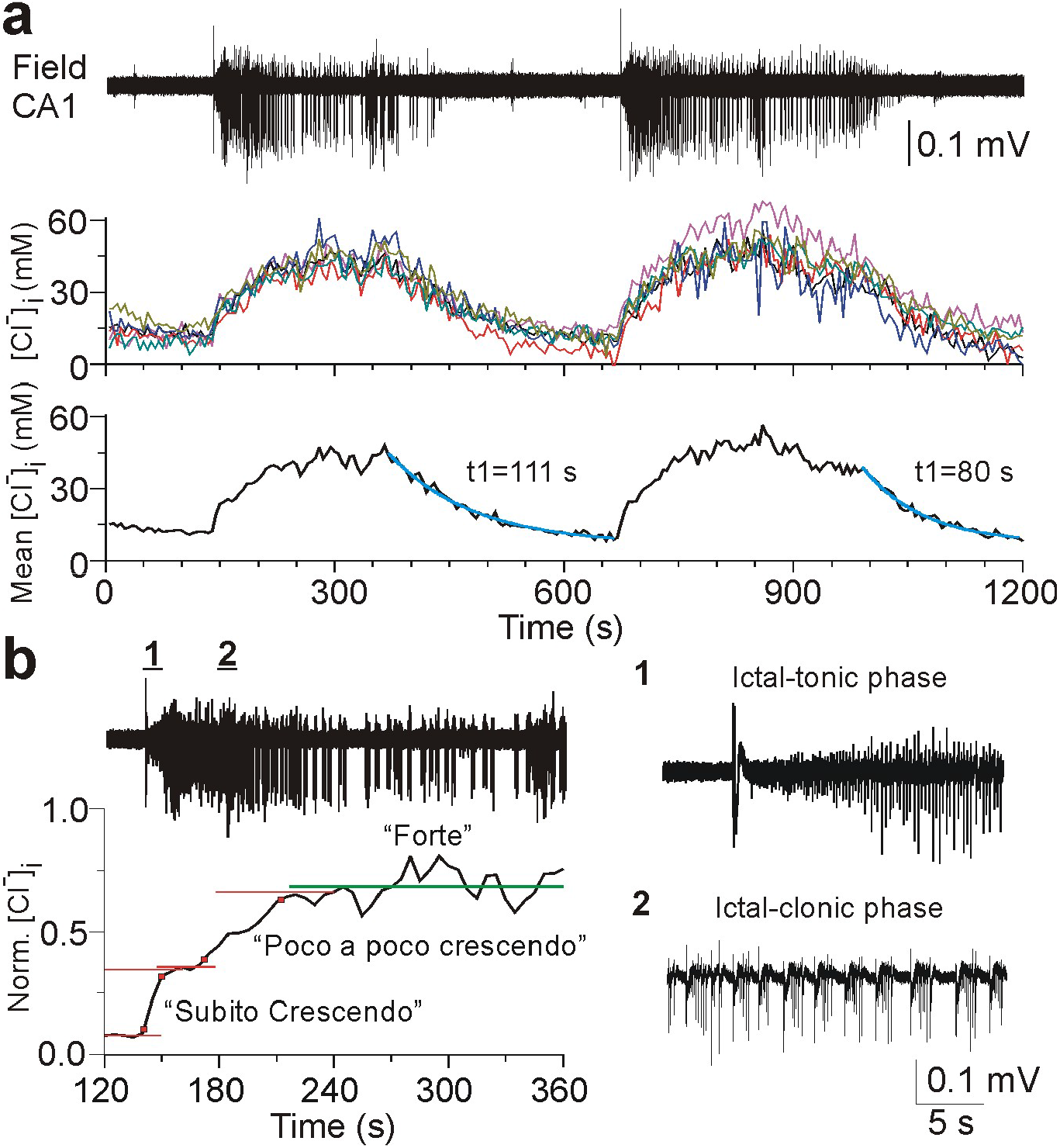
Neuronal chloride transients during spontaneous ictal-like discharges. (**a**) Simultaneous extracellular field potential recording and two-photon fluorescence chloride imaging in the organotypic hippocampal slice at DIV18. Example of spontaneous recurrent ictal-like epileptiform discharges (ILDs) and corresponding [Cl^-^]_i_ transients in the CA1 pyramidal cells (n=6), and the mean [Cl^-^]_i_. [Cl^-^]_i_ progressively increased during “crescendo” phase of sustained ictal-tonic discharges, was relatively stable during “forte” phase of intermittent ictal-clonic discharges, and progressively decreased during “decrescendo” phase of secondary after-discharges and post-ictal depression (decay time constant (t1) 80-111 s). (**b**) Expansion of ILDs and mean [Cl^-^]_i_ changes normalized to values between 0 and 1 ([0, 1]). [Cl^-^]_i_ rapidly increased during hyper-synchronous onset of ILD (“subito crescendo”) and subsequent sustained ictal-tonic discharges (“poco a poco crescendo”) and was relatively stable during intermittent epileptiform discharges. (**b1-2**) Expansion of sustained ictal-tonic discharges and intermittent ictal-clonic epileptiform discharges.

Progressive [Cl^−^]_i_ elevations during the onset of ILDs due to intensive GABAergic inputs in the pyramidal cells may shift the effects of post-synaptic GABA_A_ receptor (GABA_A_-R) mediated currents from depolarizing to excitatory and/or reverse the net effects of GABA from inhibitory to excitatory (Lillis et al., 2012;Jedlicka et al., 2011). Such changes may shift a neuronal network into a periodic bursting mode associated with sustained ictal-like discharges and intermittent after-discharges (Figure 1). Therefore, prevention of [Cl^−^]_i_ elevations and/or facilitation of [Cl^−^]_i_ extrusion may improve chloride homeostasis, GABAergic inhibition, and control of electrographic seizures.

### The WNK-SPAK/OSR1 inhibitor WNK463 enhanced neuronal chloride extrusion

The Cl^-^-sensitive WNK-SPAK/OSR1-CCCs pathway, involving WNK (with no lysine or lysine-deficient protein kinase) and downstream SPAK (SPS1-related proline/alanine-rich kinase) and OSR1 (oxidative stress-responsive kinase-1) complex, and cation-chloride cotransporters, regulates cell volume, neuronal chloride homeostasis and corresponding GABA_A_ signaling (Shekarabi et al., 2017). We determined the role of WNK-SPAK/OSR1 kinase activity in modulation of [Cl^−^]_i_ transients induced by intensive GABA_A_-R activation in principal neurons. A puff micropipette filled with extracellular solution containing GABA (50 µM) to activate GABA_A_-Rs and L-Glutamate (50 µM) to depolarize the membrane potential and maximize GABA_A_-R mediated Cl^-^ influx was placed above the dendritic area of the neurons. Focal puff application of GABA/L-Glu (puff duration 5-10 ms, pressure 5-10 psi) in ACSF containing TTX (1 µM) induced [Cl^−^]_i_ transients in a subpopulation of superficial neurons (Figure 2a). The decay time constant of baseline [Cl^−^]_i_ recovery was plotted as a function of baseline [Cl^−^]_i_. Linear regression analysis suggested a minimal interaction between baseline [Cl^-^]_i_ and the rate of [Cl^-^]_i_ recovery in individual cells that responded to GABA/L-Glu puff applications (Figure 2b; y=A+B*x, A=57.97±19.4, B=-0.26±0.87; n=6 slices, 17 cells; Pearson’s r=-0.078, R^2^=0.006).

**Figure 2.**
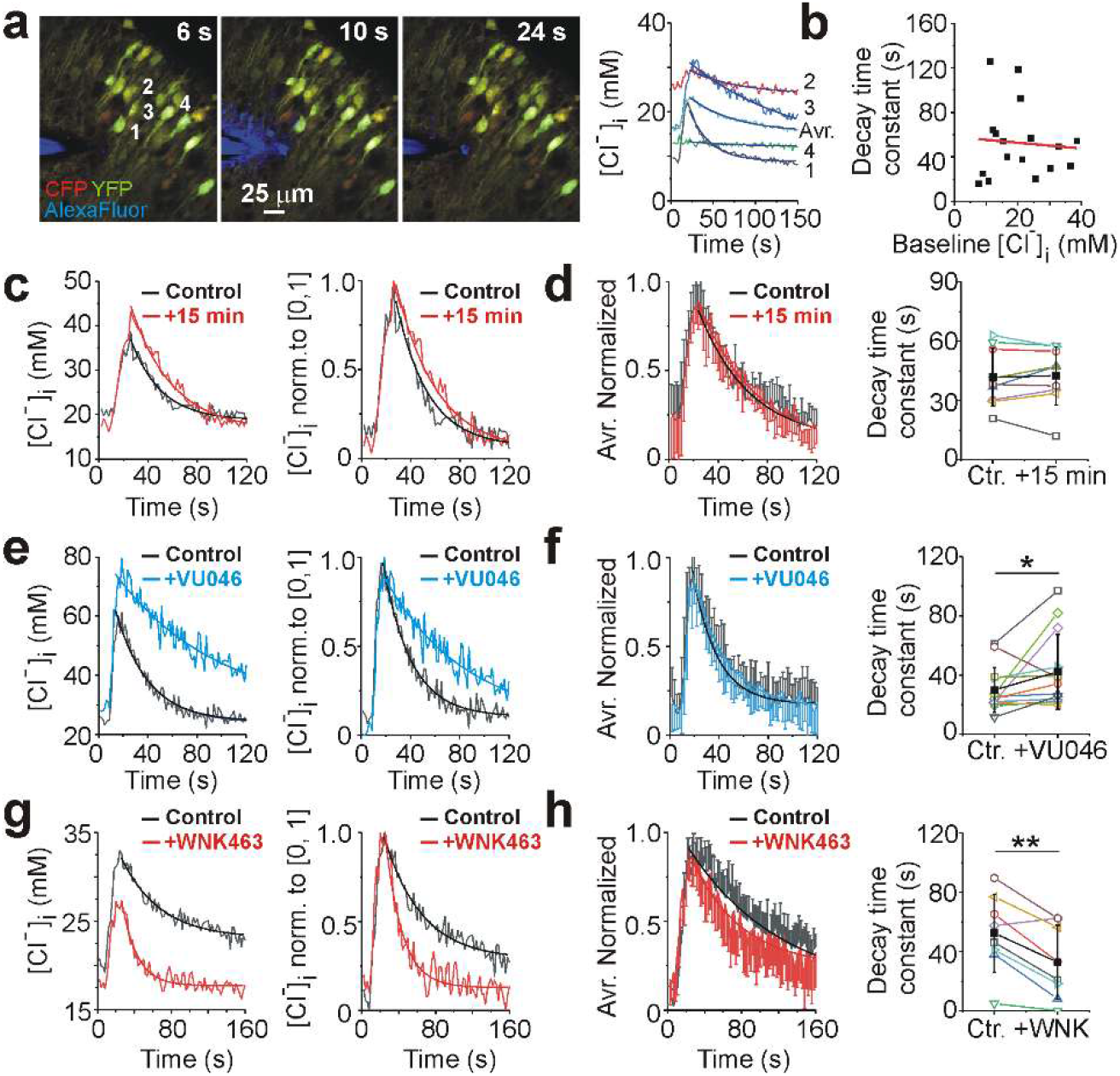
WNK-SPAK/OSR1 inhibitor WNK463 facilitated neuronal chloride extrusion. (**a**) An illustration of the designed experiment to induce neuronal chloride transients in the CA1 pyramidal cells expressing sCLM. The puff micropipette filled with extracellular solution containing GABA (50 µM) and L-Glu (50 µM) was placed above the dendritic area of the neurons. Merged CFP (*red*) and YFP (*green*) fluorescent signals in subpopulation of neurons before (6 s), during (10 s) and after (24 s) GABA/L-Glu puff application. Alexa Fluo-594 (*blue*) was added for visualization. Corresponding [Cl^−^]_i_ transients induced by GABA/L-Glu puff application. Exponential decay fit (y=y_0_+A1exp(-(x-x_0_)/t1)) was used to measure decay time constant (t1) of [Cl^-^]i recovery in individual cells (1-4; *solid curves*). (**b**) Linear regression analysis revealed that decay time constant of [Cl^-^]_i_ recovery in individual cells was independent of the baseline [Cl^-^]_i_ and corresponding E_GABA_. (**c-h**) [Cl^−^]_i_ transients, corresponding [Cl^-^]_i_ changes normalized to values between 0 and 1 and exponential decay curves in individual cells and averaged means (mean±SD) in control (**c, d**), before and during 1 µM VU0463271 application (**e, f**) and 1 µM WNK463 application (**g, h**). (**d**) In control, exponential decay time constant was not significantly different between two consecutive Cl^-^ transients. (**f**) The KCC2 antagonist VU0463271 significantly increased the mean decay time constant (*P<0.05, paired t-test). (**h**) The WNK-SPAC/OSR1 inhibitor WNK463 significantly decreased the mean decay time constant of chloride recovery (**P<0.01, paired t-test).

The decay time constant of [Cl^−^]_i_ recovery in individual cells was compared in control conditions, before and during application of the WNK-SPAK/OSR1 inhibitor WNK463 (1 µM), and the KCC2 inhibitor VU04663271 (1 µM) (Figure 2c-h). To facilitate comparisons between neurons, the chloride transients in cells that responded to GABA/L-Glu puff applications were normalized to values between 0 and 1. Under control conditions, the mean decay time constant of [Cl^-^]_i_ recovery (extrusion) was not significantly different between two consecutive (10-15 min interval) chloride transients (Figure 2c, d; n=5 slices at DIV9-14, 9 cells; 41.73±14.65 compared to 42.54±14.75 s; paired t-test, t=-0.4, P=0.69). Thus, individual neurons within the same slice showed reproducible responses to repeated GABA/L-Glu puffs. However, the variance of responses between slices was much larger, reflecting the variable geometry of the puffer pipette to the dendritic glutamate and GABA_A_ receptors.

In line with previous studies (Dzhala and Staley, 2021), bath application of the KCC2 inhibitor VU0463271 (1 µM for 10-15 min) significantly increased the mean decay time constant of baseline chloride recovery from 28.8±13.4 to 39.8±25.6 s (Figure 2e-f; n=6 slices at DIV9-12, 14 cells; paired t-test, t=-2.018, P=0.017). In contrast, bath application of WNK463 (1 µM for 15-20 min) significantly decreased the mean decay time constant from 52.5±26.2 to 32.8±24.9 s (Figure 2g-h; n=5 slices at DIV9-14, 8 of 10 cells; paired t-test, t=4.27, P=0.004). In two of ten cells, the exponential decay function did not fit the recovery of [Cl^-^]_i_ (not shown). In all cells that responded to GABA/L-Glu puff applications WNK463 also significantly reduced the amplitude of [Cl^-^]_i_ transients, which we tentatively attribute to enhanced Cl^-^ extrusion. Thus, the WNK-SPAK/ORS1 inhibitor WNK463 enhanced [Cl^-^]_i_ extrusion during recovery from GABA/L-Glu induced chloride transients.

Next, we determined the net effects of WNK-SPAK/OSR1 catalytic activity inhibitor WNK463 on the frequency and duration of recurrent interictal- and ictal-like epileptiform discharges, baseline chloride changes, [Cl^-^]_i_ elevation and extrusion rates during ILDs, and the correlation between ionic and electrographic effects in *in vitro* model of post-traumatic epileptogenesis (Figures 3-5).

**Figure 3.**
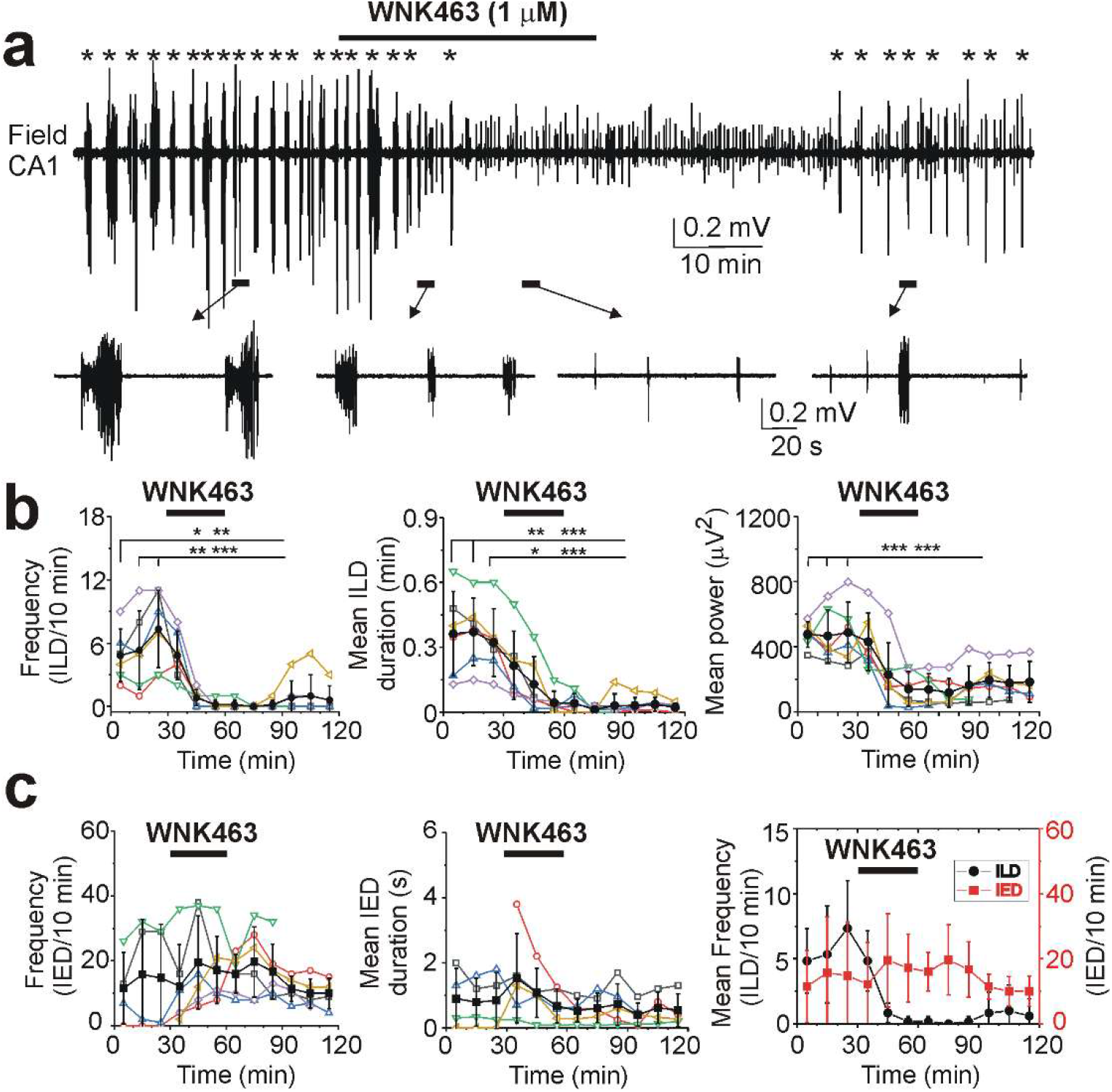
WNK-SPAK/OSR1 inhibitor WNK463 abolished recurrent ictal-like epileptiform discharges. (**a**) Extracellular field potential recording in the CA1 pyramidal cell layer in the organotypic hippocampal slice at DIV21 before (control), during and after application of 1 µM WNK463 for 30 min. Expansion of recurrent ictal (ILDs) and interictal (IEDs) epileptiform discharges before, during and after application of WNK463. (**b, c**) Corresponding summary plots of the frequency and duration of recurrent ILDs (**b**) and IEDs (**c**) in individual slice cultures (DIV14-22, open symbols) and corresponding power of electrical activity in 10 min windows. Filled symbols indicate group mean ± SD. WNK463 (1 µM) progressively decreased the mean frequency, duration, and power of recurrent ILDs and abolished ILDs (*P<0.05; **P<0.01; ***P<0.001, One Way RM ANOVA, Tukey Test). (**c**) WNK463 (1 µM) did not significantly change the mean frequency and duration of periodic IEDs (P>0.05).

### WNK463 abolished recurrent ILDs

Extracellular field potential recordings were performed in the CA1 pyramidal cell layer in the organotypic hippocampal slices at DIV12-19 before (control), during and after bath application of WNK463 (Figure 3a). Bath application of WNK463 (1 µM for 30 min) progressively reduced the mean frequency of ILDs from 7.33±3.7 to 0.17±0.41 ILD/10 min and abolished ILDs (Figure 3b; n=6 slices; One Way RM ANOVA: DF=53, F(5, 40)=15.18, P<0.001; Tukey test df=7.16, q=9.81, p<0.001). WNK463 substantially decreased the mean duration of ILDs from 0.32±0.16 to 0.04±0.05 min (DF=53, F(5, 40)=17.27, P<0.001; Tukey test: df=0.28, q=7.77, p<0.001). The corresponding mean power of electrical activity significantly decreased from 486.35±187.5 to 138.93±107.15 µV^2^ (DF=53, F(5, 40)=21.74, P<0.001; Tukey test: df=347.4, q=10.7, p<0.001). However, WNK463 did not change the mean frequency and duration of periodic IEDs (Figure 3c; IED frequency: One Way RM ANOVA, DF=47, F(5, 34)=1.01, P=0.45; IED duration: DF(5, 34)=47, F=1.41, P=0.23). Thus, WNK463 progressively reduced the frequency, duration, and power of ILDs, and abolished recurrent ILDs.

### WNK463 reduced activity-dependent interictal [Cl^-^]_i_ between ILDs

We next determined whether the anti-ictal effects of WNK463 (1 µM) correlate with changes in interictal [Cl^-^]_i_ and the rates of [Cl^-^]_i_ elevation and extrusion during recurrent ILDs (Figure 4). To clarify terminology, interictal (IED) [Cl^−^]_i_ is the cytoplasmic Cl^-^ concentration measured between ILDs. Baseline [Cl^−^]_i_ is the cytoplasmic Cl^-^ concentration measured in TTX. In line with previous studies (Lillis et al., 2012;Glykys et al., 2014;Dzhala and Staley, 2021), the median interictal [Cl^-^]_i_ transiently increased in all pyramidal cells from 25.75 (14.63-47.64) to 47.64 (25.79-88.32) mM during ILDs (Figure 4 a-c; n=4 slices at DIV12-19, 73 paired cells; Friedman RM ANOVA on Ranks, Chi-square=134.1, P<0.001; Tukey test: dr=105.6, q=9.56, p<0.05). Bath application of WNK463 abolished ILDs and corresponding [Cl^-^]_i_ transients and significantly reduced the median interictal [Cl^-^]_i_ to 23.37 (11.55-35.21) mM (dr=69.5, q=6.3, p<0.05). Interictal [Cl^-^]_i_ recovered to 26.06 (13.76-39.34) mM during washing out of WNK463 (dr=49.5, q=4.49, p<0.05). WNK463 induced changes in interictal [Cl^-^]_i_ were plotted as a function of initial interictal [Cl^-^]_i_ (Figure 4d). Linear regression analysis suggested a larger effect of WNK463 in neurons with higher initial interictal [Cl^-^]_i_ (A=11.2±1.4, B=-0.7±0.03; Pearson’s r=-0.94, R^2^=0.89). Thus, pharmacological inhibition of the WNK-SPAK/OSR1 catalytic activity by WNK463 leads to reduction in the interictal [Cl^-^]_i_ in neurons and negative shifts in the interictal E_GABA_, increasing the net inhibition of neuronal network activity, and suppression of recurrent ILDs.

**Figure 4.**
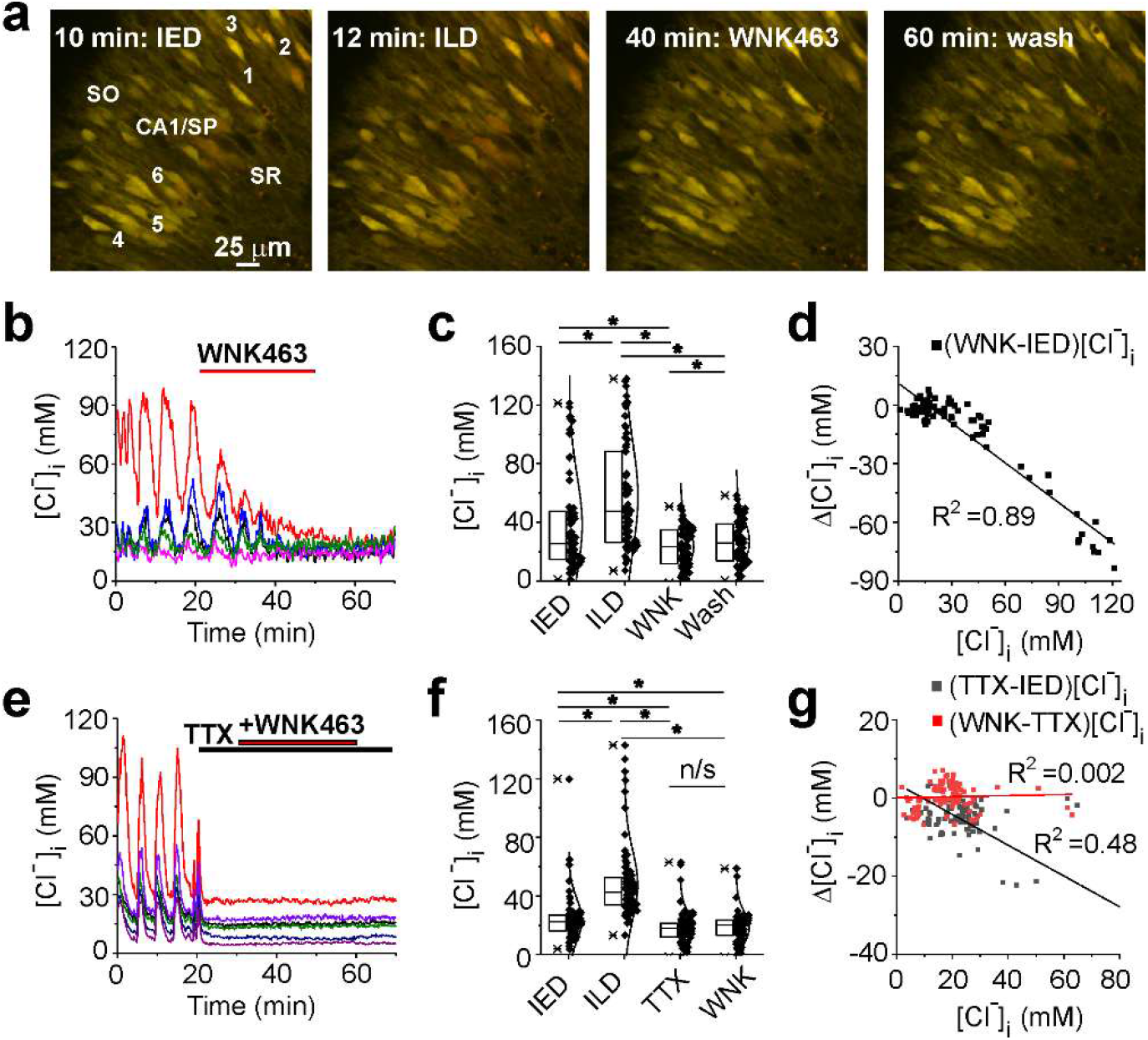
WNK463 progressively reduced activity-induced elevations in interictal [Cl^-^]_i_ and abolished neuronal Cl^-^ transients. (**a**) Two-photon microscopy images of Clomeleon in the CA1 pyramidal cell layer in the organotypic hippocampal slice *in vitro* (DIV19). Merged CFP (*red*) and YFP (*green*) fluorescent signals before, during and after WN463 (1 µM for 30 min) application. (**b**) Corresponding [Cl^-^]_i_ changes in individual cells are plotted as a function of time. WNK463 abolished recurrent ILDs and corresponding Cl^-^ transients. (**c**) Baseline [Cl^−^]_i_ distribution in subpopulations of neurons before (IED and ILD phases), during and after application of WNK463. Box (*left*) + data (*right*) plots correspond to median (25%–75%) [Cl^−^]_i_ in individual cells (filled symbols) and their Gaussian distribution curves. WNK463 significantly reduced the median interictal [Cl^−^]_i_ (*P<0.05; Friedman RM ANOVA on Ranks). (**d**) Corresponding interictal [Cl^−^]_i_ changes induced by WNK463. Data were fitted with a linier regression fit. (**e**) WNK463 (1 µM) did not change baseline [Cl^-^]_i_ when synaptic activity had already been suppressed with the sodium channel blocker TTX (1 µM). (**f**) Baseline [Cl^−^]_i_ distribution and corresponding [Cl^−^]_i_ changes induced by TTX and WNK463 in the presence of TTX (*P<0.05; Friedman RM ANOVA on Ranks). (**g**) Corresponding interictal [Cl^−^]_i_ changes in individual cells induced by TTX (black symbols and line) and baseline [Cl^−^]_i_ changes induced by WNK463 in the presence of TTX (red symbols and line). Data were fitted with a linear regression fit.

**Figure 5.**
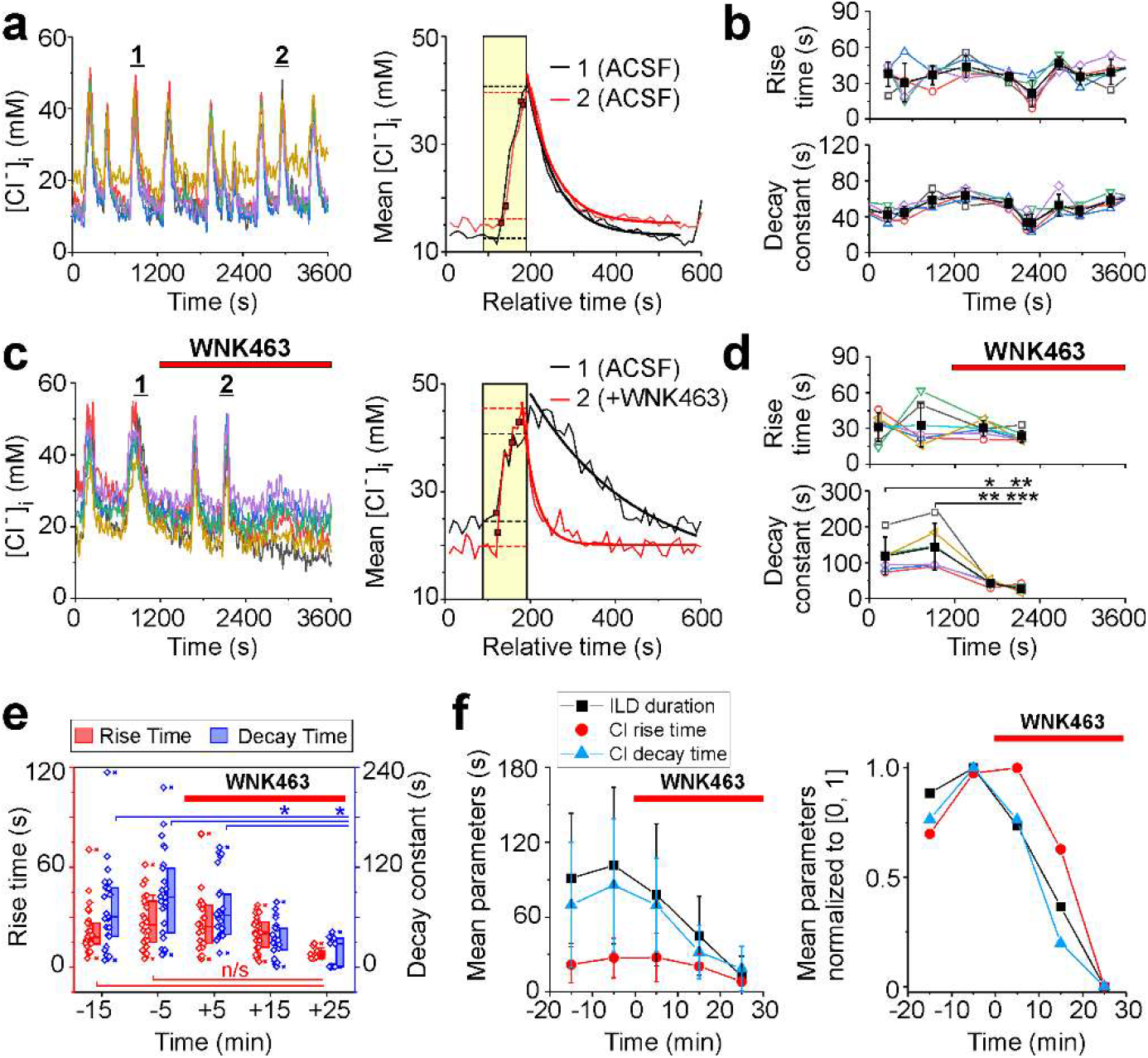
Effects of the WNK-SPAK/OSR1 blocker WNK463 on neuronal chloride transients during recurrent ictal-like epileptiform discharges. (**a, c**) Neuronal Cl^-^ changes in individual cells as a function of time and corresponding mean [Cl^−^]_i_ transients in control ACSF (**a1-2**), and before and during WNK463 (1 µM) application (**c1-2**). (**b, d**) Corresponding rise time and decay time constant of [Cl^−^]_i_ transients in individual cells (open symbols) as a function of time in control (**b**), and before and during WNK463 application (**d**). Filled symbols indicate group mean ± SD. WNK463 progressively decreased the mean decay time constant of [Cl^−^]_i_ transients (*P<0.05, **P<0.01, ***P<0.001; One Way RM ANOVA). (**e**)[Cl^−^]_i_ rise time and decay time constant of neuronal chloride transients in individual cells before and during WNK463 application (n/s corresponds to P>0.05, *P<0.05, Friedman RM ANOVA on Ranks, Tukey test). Box (*right*) + data (*left*) plots correspond to median (25%–75%) [Cl^−^]_i_ in individual cells (open symbols). (**f**) The mean ILD duration, corresponding [Cl^-^]_i_ rise time and decay time constant of neuronal chloride transients, and corresponding normalized parameters before and during WNK463 application. WNK463 progressively reduced the mean duration of ILDs (black symbols) in line with enhanced Cl^-^ extrusion rate (blue symbols).

Pharmacological manipulations of the WNK-SPAK/OSR1-CCCs activity by WNK463 could produce anti-ictal effects that affect [Cl^-^]_i_ as a consequence of reduced ictal Cl^-^ loading, separately from the consequences of altered chloride transport (Dzhala and Staley, 2021). To estimate how direct anti-ictal effects might alter interictal [Cl^-^]_i_, we applied the sodium channel antagonist TTX (1 μM) to block synaptic activity and determined the effect of WNK463 on baseline [Cl^-^]_i_ in the presence of TTX (Figure 4e-g). TTX application rapidly blocks ILD activity directly, providing a means to assess the effects of the intensity of ILD activity on interictal [Cl^-^]_i_ levels separately from effects on Cl^-^ transport. TTX rapidly abolished ILDs and corresponding [Cl^-^]_i_ transients, and significantly reduced the median interictal [Cl^-^]_i_ from 22.5(16.06-26.94) to a baseline of 17.98 (11.94-21.36) mM (Figure 4e-f; N=4 slices at DIV14-16, 90 paired cells; Friedman RM ANOVA on Ranks, Chi-square=206.17, P<0.001; Tukey test: interictal [Cl^-^]_i_ compared to TTX, dr=111.5, q=9.1, p<0.05). Subsequent application of WNK463 (1 µM) in the continued presence of TTX did not significantly change the median baseline [Cl^-^]_i_ to 20.25(13.16-23.4) mM ( dr=11.5, q=0.94, p>0.05) suggesting a contribution of activity-dependent [Cl^-^]_i_ accumulation during recurrent ILDs. However, the median baseline [Cl^-^]_i_ in the presence of WNK463+TTX remained significantly lower compared to the no-TTX control interictal condition (dr=100, q=8.16, p<0.05). TTX induced changes in interictal [Cl^-^]_i_ were plotted as a function of initial interictal [Cl^-^]_i_, and WNK463 induced changes in baseline [Cl^-^]_i_ were plotted as a function of preceding baseline [Cl^-^]_i_ in the presence of TTX (Figure 4g). Linear regression analysis revealed that the rate of interictal [Cl^-^]_i_ reduction induced by TTX was greater than the effect of WNK463 on baseline [Cl^-^]_i_ in the presence of TTX (Figure 4g: Δ[Cl^-^]_i_ (TTX) vs interictal [Cl^-^]_i_ in control: A=3.68±1.2, B=-0.39±0.04, Pearson’s r=-0.69, R^2^=0.48; Δ[Cl^-^]_i_ (WNK463+TTX) vs baseline [Cl^-^]_i_ in the presence of TTX: A=0.026±0.7, B=0.014±0.03, Pearson’s r=0.043, R^2^=0.002). Thus, pharmacological inhibition of the WNK-SPAK/OSR1 catalytic activity by WNK463 leads to both increased CCC-mediated Cl^-^ export as well as indirect decreases in Cl^-^ influx during ictal network activity that are replicated by TTX. Both effects increase the net inhibition of neuronal network activity and suppression of recurrent ILDs.

### WNK463 progressively enhanced neuronal chloride extrusion during ILDs

As another assessment of the effects of WNK-SPAK/OSR1 inhibition on neuronal chloride transport activity during suppression of recurrent ILDs, we compared the rise-time of [Cl^-^]_i_ elevation and decay time constant of [Cl^-^]_i_ extrusion during recurrent ILDs in control, and before and during application of WNK463 (1 µM) (Figure 5a, c). In control conditions, the mean rise time of [Cl^-^]_i_ elevation and the mean decay time constant of [Cl^-^]_i_ extrusion during recurrent ILDs were relatively stable in line with the frequency and duration of recurrent ILDs (n=5 slices at DIV13-18, n=30 paired cells; Rise time: One Way RM ANOVA, DF=89, F(29, 58)=0.76, P=0.471; Decay time constant: DF=89, F(29,58)=1.26, P=0.292). WNK463 application progressively depressed recurrent ILDs (Figure 3), providing a means to assess the effects of WNK463 on neuronal [Cl^-^]_i_ transients and kinetics. Bath application of WNK463 (1 µM) did not significantly reduce the median rise time of chloride accumulation during onset of ILDs from 25.5(14.3-40.3) s in control, before WNK463 application, to 20.7(11.8-27.8) s over 10 to 20 min WNK463 application (Figure 5e; n=5 slices at DIV12-19, 25 paired cells; Friedman RM ANOVA on Ranks, Chi-square=15.44 with 4 degrees of freedom, P=0.004; Tukey test: dr=4, q=0.7, p>0.05). Recurrent ILDs and [Cl^-^]_i_ transients were suppressed over 20 to 30 min WNK463 application (Figures 3, 5). The corresponding exponential decay time constant (t1) of [Cl^-^]_i_ extrusion (recovery) during ILDs significantly decreased from the median 83.9(39.7-119.7) s in control to 30(17.8-46.5) s in the presence of WNK463 (Figure 5e; Friedman RM ANOVA on Ranks, Chi-square=31.9 with 4 degrees of freedom, P<0.001; Tukey test: dr=31.5, q=5.52, p<0.05). In contrast, application of KCC2 inhibitor VU0463271 (1 µM) increased the decay time constant of [Cl^-^]_i_ extrusion during recurrent ILDs and corresponding chloride transients in line with increased duration of recurrent ILDs and transition to electrical *status epilepticus* (Dzhala and Staley, 2021). Thus, the WNK-SPAK/OSR1 pathway can regulate the maximum velocity of CCCs during intense activation of the GABA_A_-R and [Cl^-^]_i_ accumulation as occurs during prolonged ILDs. Progressively decreased duration and power of recurrent ILDs in the presence of the WNK-SPAK/OSR1 blocker WNK463 would be expected to decrease Cl^-^ influx, and correlates with decreased rise time, amplitude, and duration of [Cl^-^]_i_ elevation, consistent with both a decrease in Cl^-^ influx and an enhanced extrusion rate of [Cl^-^]_i_ (Figure 5f).

### WNK463 had no effects in the presence of the GABA_A_-receptor antagonist and CCCs inhibitors

We next used pharmacological tools to determine whether GABA_A_-R block prevents the anti-ictal effects of the WNK-SPAK/OSR1 blocker WNK463 (Figure 6a-c). In line with previous studies (Dzhala and Staley, 2021), bath application of the GABA_A_-R antagonist SR95531 (10 µM) rapidly abolished ILDs and corresponding [Cl^-^]_i_ transients (Lillis et al., 2012), and induced large amplitude and 1-2 s duration interictal-like epileptiform discharges (IEDs) (Figure 6a, b; n=6 slices at DIV13 to DIV17, ILD frequency: One Way RM ANOVA, DF=71, F(5, 55)=12.53, P<0.001). Subsequent application of WNK463 (1 µM), in the continued presence of SR95531, did not significantly change the mean frequency and duration of epileptiform discharges, and the corresponding mean power of electrical activity in 10 min windows (Figure 6a-c; IED frequency: One Way RM ANOVA, DF=58, F(4, 43)=1.78, P=0.088; Power: One Way RM ANOVA, DF=71, F(5, 55)=1.346, P=0.225). These data suggest that the anti-ictal effects of the WNK-SPAK/OSR1 blocker WNK463 require activation of GABA_A_-Rs. However, we cannot rule out that cation-chloride cotransporter (CCC)-mediated Cl^-^ export and GABA_A_-R activity is not as critical to the termination of short-duration IEDs (Staley et al., 1998) as to the termination of longer-lasting ILDs.

**Figure 6.**
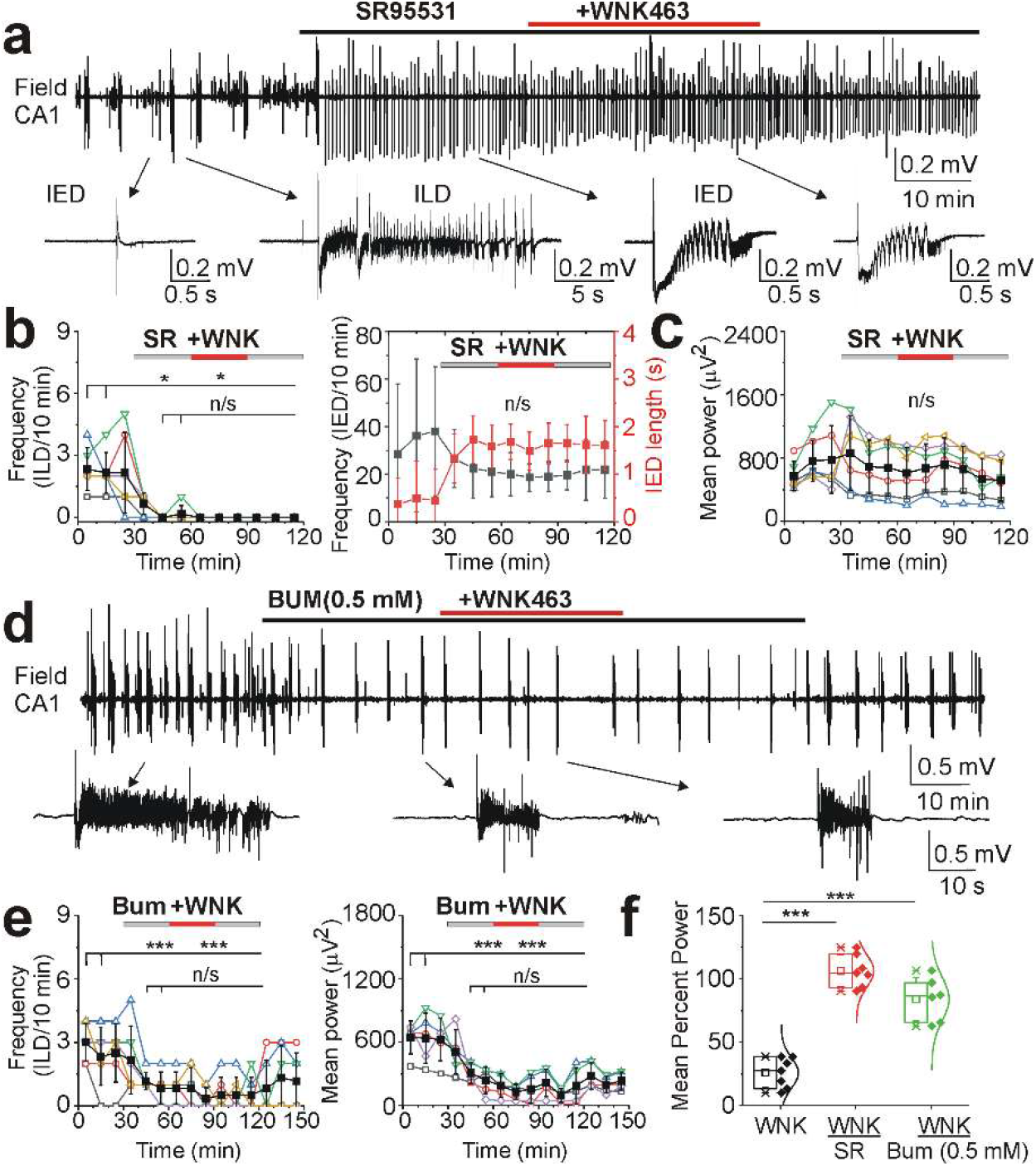
WNK463 had no effects in the presence of the GABA_A_-receptor antagonist and CCCs inhibitors. (**a, d**) Extracellular field potential recordings in the CA1 pyramidal cell layer in the organotypic hippocampal slices *in vitro*. Expansion of recurrent interictal (IEDs) and ictal (ILDs) epileptiform discharges in control and during drug applications. (**b, c**) Application of the GABA_A_-R antagonist SR95531 (10 µM) abolished spontaneous ILDs and induced large amplitude IEDs. Subsequent application of WNK463 (1 µM) in the presence of SR95531 did not change the mean frequency and duration of epileptiform discharges and corresponding power of electrical activity (n/s – P>0.05; *P<0.05; Friedman RM ANOVA on Ranks, Dunn’s test). (**e**) Cation-chloride cotransporter blocker bumetanide (0.5 mM) reduced the mean frequency of ILDs, corresponding power of electrical activity, and prevented the anti-ictal effects of WNK463 ( n/s – P>0.05; *P<0.05; ***P<0.001; One Way RM ANOVA, Tukey test). (**f**) The mean effect of WNK463 on the power of epileptiform activity (percent of treatment-preceding power) was significantly different as the corresponding effects of WNK463 in the presence of 10 µM SR95531 or 0.5 mM bumetanide (***P<0.001; One Way ANOVA, Holm-Sidak Test).

High concentrations of CCC inhibitors have large anticonvulsant effects (Hochman et al., 1995;Dzhala and Staley, 2021). We also evaluated whether CCCs were necessary for the anti-ictal effects of WNK463 (Figure 6d-f). Under similar experimental conditions (DIV12 to DIV16), bath application of a high concentration of bumetanide (0.5 mM) that nonspecifically blocks CCCs (NKCC1 and KCC2-3) (Payne, 1997) significantly reduced the mean frequency of spontaneous ILDs from 3±0.89 to 0.83±0.75 ILD/10 min. Subsequent application of WNK463 (1 µM) in the continued presence of 0.5 mM bumetanide did not significantly change the mean frequency of ILDs to 0.5±0.84 ILD/10 min (Figure 6e; N=6 slices; One Way RM ANOVA: DF=71, F(5, 55)=9.71, P<0.001; Tukey Test: control compared to bumetanide, df=2.17, q=7.2, P<0.001; control compared to WNK463 in the presence of bumetanide, df=2.5, q=8.3, P<0.001; bumetanide compared to WNK463 in the presence of bumetanide, df=0.33, q=1.1, P=1.00). The corresponding mean power of electrical activity significantly decreased from 644±153.97 µV^2^ in control to 238.49±127.18 µV^2^ during bumetanide application, and consecutive application of WNK463 in the presence of bumetanide did not significantly change the mean power of electrical activity to 215.3±128.9 µV^2^ (Figure 6e; One Way RM ANOVA: DF=59, F(4, 44)=15.16, P<0.001; Tukey Test: control compared to bumetanide, df=405.53, q=7.4, P<0.001; control compared to WNK463 in the presence of bumetanide, df=428.69, q=7.8, P<0.001; bumetanide compared to WNK463 in the presence of bumetanide, df=23.16, q=0.42, P=0.987).

The mean effect of WNK463 on the power of epileptiform activity (percent of treatment-preceding power) was significantly different as the corresponding effects of WNK463 in the presence of 10 µM SR95531 or 0.5 mM bumetanide (Figure 6f; One Way ANOVA, DF=19, F(2, 17)=62.2, P<0.001; Holm-Sidak Test: WNK463+SR95531 compared to WNK463, df=80.47, t=10.83, P<0.001; WNK463+Bum compared to WNK463, df=58.27, t=7.53, P<0.001). These results suggest that the anti-ictal action of WNK463 requires functional GABA_A_-Rs and is dependent on CCCs activity, and greatly exceeds the anti-ictal effects of specific blockers of NKCC1 transport activity or activators of KCC2 transport activity under similar experimental conditions (Dzhala and Staley, 2015;Dzhala and Staley, 2021). However, the WNK effect is in line with the anticonvulsant effects of very high, non-specific concentrations of CCC antagonists (Hochman et al., 1995;Dzhala and Staley, 2021).

### NKCC1 inhibition did not prevent the anti-ictal effects of WNK463

The WNK-SPAK/OSR1 catalytic pathway inhibitor WNK463 reduces phosphorylation of the WNK1 downstream targets SPAK/OSR1 (Yamada et al., 2017) and thereby inhibits NKCC1 and KCC2 phosphorylation (de Los et al., 2014), resulting in inhibition of NKCC1 and stimulation of KCC2 activity. The relative contribution of NKCC1 and KCC2 transport activity to the anti-ictal effects of WNK463 is not known. We used pharmacological tools to determine whether NKCC1 inhibition by bumetanide (10 μM) and KCC2 inhibition by VU0463271 (1 µM) prevents the anti-ictal effects of WNK463 (Figure 7).

**Figure 7.**
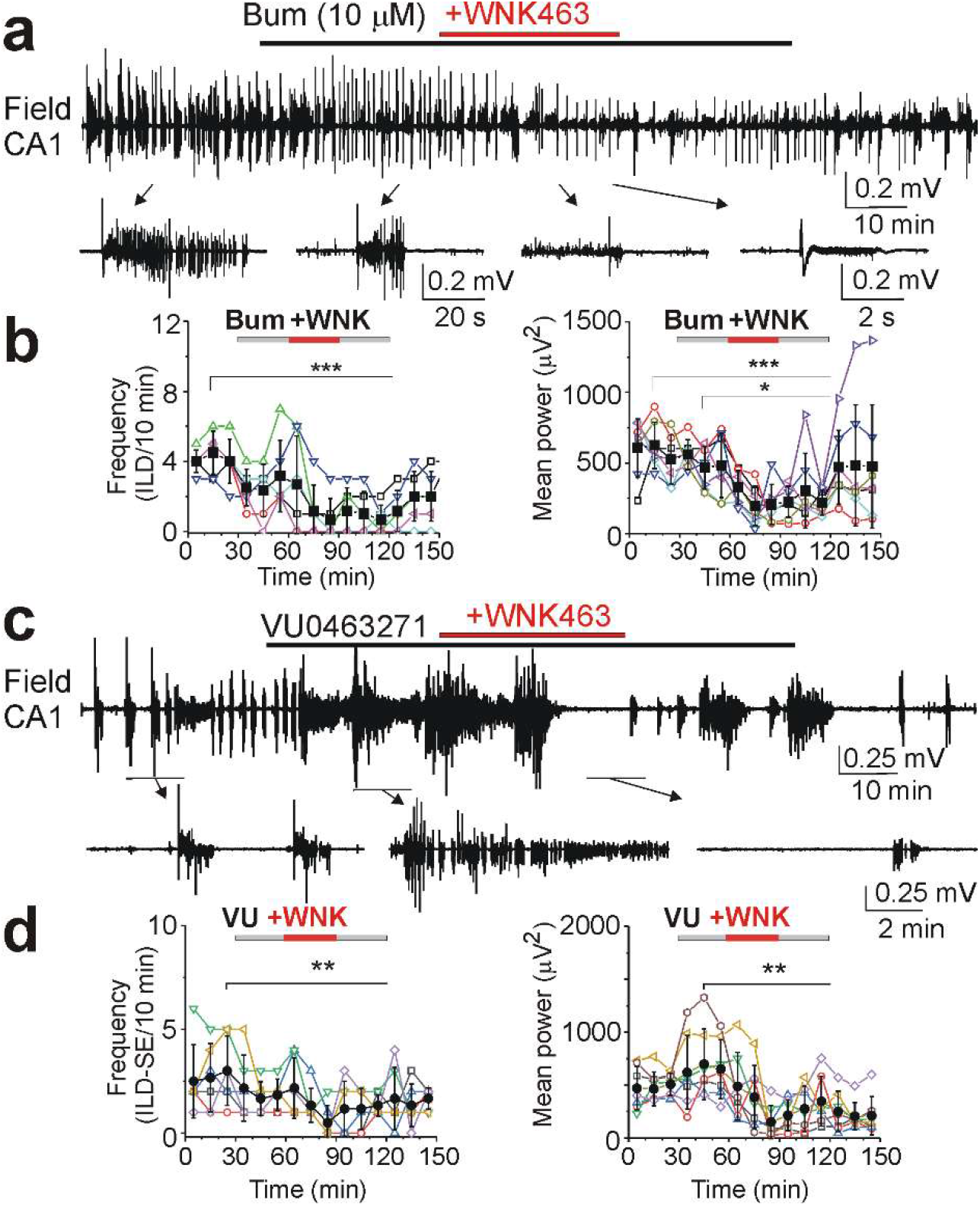
Anti-ictal effects of WNK463 in the presence of NKCC1 blocker bumetanide and KCC2 blocker VU0463271. (**a, c**) Extracellular field potential recordings in the CA1 pyramidal cell layer in the organotypic hippocampal slices *in vitro*. WNK463 (1 µM) was applied in the presence of (**a**) NKCC1 blocker bumetanide (10 µM) and (**c**) KCC2 blocker VU0463271 (1 µM). Expansion of epileptiform discharges before and during drugs applications. (**b**) WNK463 application in the presence of bumetanide significantly reduced the mean frequency of recurrent ILDs in control and the mean power of electrical activity. (**d**) WNK463 in the presence of VU0463271 significantly reduced the mean frequency of ILDs in control and the mean power of electrical activity in the presence of VU0463271 alone (*p<0.05, **p<0.01, ***p<0.001; One-Way RM ANOVA, Tukey test).

Bath application of bumetanide (10 µM) did not significantly change the mean frequency of recurrent ILDs from 4.5±1.2 to 3.2±2.04 ILD/10 min, and subsequent application of WNK463 (1 µM) in the continued presence of bumetanide (10 µM) reduced the mean frequency of ILDs to 0.67±1.2 ILD/10 min (Figure 7b; n=6 slices at DIV16 to 21; One Way RM ANOVA: DF=71, F(5,55)=7.1, P<0.001; Tukey Test: control compared to bumetanide, df=1.33, q=2.51, p=0.75; control compared to WNK463+bumetanide, df=3.83, q=7.16, p<0.001; bumetanide compared to WNK463+bumetanide, df=2.5, q=4.7, p=0.05). Bumetanide (10 µM) did not significantly change the mean power of corresponding electrical activity from 624.8±164.5 to 467.71±149.83 µV^2^, and consecutive application of WNK463 in the presence of bumetanide significantly decreased the mean power of electrical activity to 194.84±138.94 µV^2^ (Figure 7b; One Way RM ANOVA: DF=71, F(5, 55)=5.48, P<0.001; Tukey Test: control compared to bumetanide, df=157.11, q=2.72, p=0.59; control compared to WNK463+bumetanide, df=429.9, q=7.46, p<0.001; bumetanide compared to WNK463+bumetanide, df=272.87, q=4.73, p=0.039). On average, WNK463 application in the presence of bumetanide significantly reduced the mean power of electrical activity in the presence of bumetanide (10 µM) alone by 59.9±21.1%. These data indicate that the effect of WNK463 is not mediated solely by effects on NKCC1 activity.

### KCC2 inhibition did not prevent the anti-ictal effects of WNK463

We next determined whether KCC2 inhibition affects the anti-ictal effects of WNK463 (Figure 7c). Under similar experimental conditions (DIV16 to DIV21), bath application of KCC2 inhibitor VU0463271 (1 µM) reduced the mean frequency of ILDs from 3±1.67 to 1.83±0.75 ILD/10 min, and subsequent application of WNK463 (1 µM), in the continued presence of VU0463271, reduced the mean frequency of ILDs to 0.5±0.84 ILD/10 min (Figure 7d; n=6 slices, One Way RM ANOVA: DF=71, F(5, 55)=3.46, P=0.001; Tukey Test: control compared to VU0463271, df=1.17, q=2.86, p=0.59; control compared to WNK463+VU0463271, df=2.5, q=6.13, p=0.003; VU0463271 compared to WNK463+ VU0463271, df=1.33, q=3.27, p=0.4). VU0463271 increased the mean power of corresponding electrical activity from 498.37±148.11 µV^2^ in control to 697.55±335.23 µV^2^, and consecutive application of WNK463, in the presence of VU0463271, significantly decreased the mean power of electrical activity to 151.67±157.47 µV^2^ (Figure 7d; One Way RM ANOVA: DF=71, F(5,55)=3.03, P=0.003; Tukey Test: VU0463271 compared to control, df=199.18, q=2.38, p=0.86; control compared to WNK463+VU0463271, df=346.7, q=4.15, p=0.15; VU0463271 compared to WNK463+VU0463271, df=545.9, q=6.53, p=0.001). On average, WNK463 application in the presence of VU0463271 reduced the mean power of electrical activity in the presence of VU0463271 alone by 44.8±32.3%. These results indicate that the anti-ictal effects of WNK463 are not mediated solely by effects on KCC2 activity and are likely due to coherent inhibition of NKCC1 and activation of KCC2.

### siRNA targeted Slc12a2 or Slc12a5 gene silencing did not prevent the anti-ictal effects of WNK463

We next determined whether small-interfering RNA (siRNA) targeted gene silencing of either Slc12a2 (NKCC1) or Slc12a5 (KCC2) individually prevent or reduce the anti-ictal effects of WNK463, in line with pharmacological inhibition of NKCC1 and KCC2. To assess uptake of siRNA, age-matched organotypic hippocampal slices were prepared, cultured and incubated in corresponding siRNA delivery medium. After 72 hours of incubation with green (fluorescein) non-targeting siRNA (DIV9-11), uptake of siRNA was visible during an additional 24+ hours incubation in standard culture medium (Figure 8a). The most prominent and intense fluorescence was observed in cells of *stratum radiatum* and *stratum pyramidale* layer, indicating an uptake of siRNA. Extracellular field potential recording in the CA1 and CA3 pyramidal cell layer in the siRNA-treated organotypic hippocampal slices revealed spontaneous multiple unit activity and population neuronal network activity, including short interictal-like discharges and prolonged ictal-like discharges (Figure 8b).

**Figure 8.**
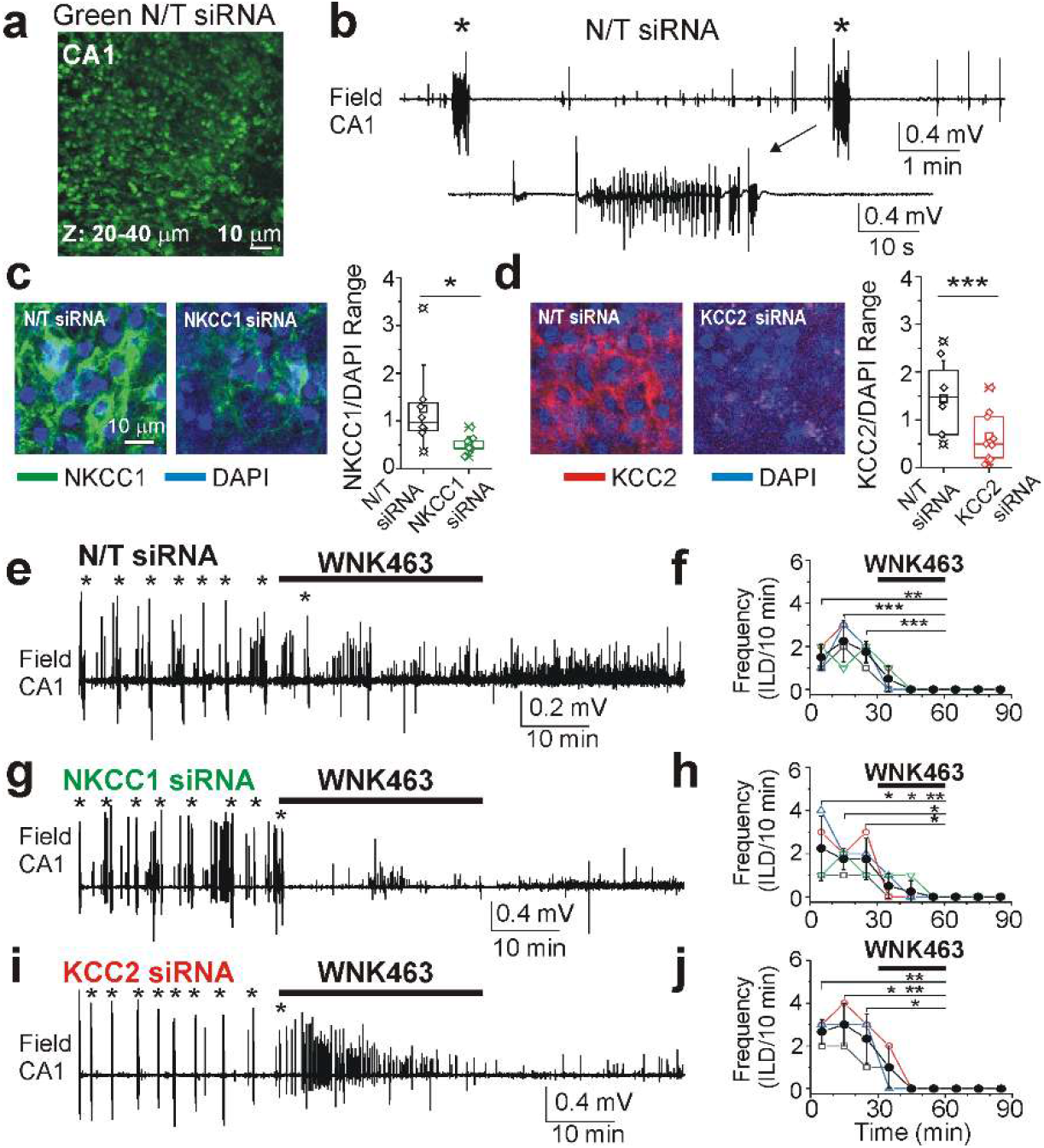
siRNA targeted Slc12A2 (NKCC1) and Slc12a5 (KCC2) gene silencing did not prevent the anti-ictal effects of WNK463. (**a**) Cellular penetration by Accell green (fluorescein) non-targeting siRNA in the organotypic hippocampal slice *in vitro* at DIV12. (**b**) Simultaneous extracellular field potential recordings in the CA1 pyramidal cell layer revealed spontaneous multiple unit activity (MUA), interictal epileptiform discharges (IEDs) and ictal-like epileptiform discharges (ILDs). Expansion of recurrent ILD during recordings. (**c, d**) NKCC1 (green) and KCC2 (red) antibodies overlaid with 4′,6-diamidino-2-phenylindole (DAPI) staining (blue) in control (N/T siRNA) and NKCC1 and KCC2 siRNA treated organotypic hippocampal slices. Optical density of NKCC1 and KCC2 immunostaining relative to DAPI staining revealed significant siRNA targeted Slc12A2 and Slc12a5 gene silencing and protein expression (*P<0.05; ***P<0.001; Mann-Whitney test on ranks). (**e, f**) WNK463 (1 µM) abolished spontaneous ILDs (marked by asterisks) in control slices treated with non-targeting (N/T) siRNA. (**g, h**) WNK463 (1 µM) abolished spontaneous ILDs (marked by asterisks) in slices treated with Slc12a2 (NKCC1) siRNA. (**i, j**) WNK463 (1 µM) abolished spontaneous ILDs (marked by asterisks) in slices treated with Slc12a5 (KCC2) siRNA (*P<0.05, **P<0.01, ***P<0.01, One-Way RM ANOVA, Tukey test).

Previously published data revealed that siRNA targeted Slc12a2 gene silencing reduced NKCC1 expression and limited injury-induced changes in neuronal volume and chloride (Bahari et al., 2024). In all three age-matched groups of organotypic hippocampal slices cultured and treated for 72 hours (DIV9-11) with non-targeting siRNA (control), Slc12a2 siRNA (NKCC1) and Slc12a5 siRNA (KCC2), immunohistochemistry using specific antibodies revealed NKCC1 and KCC2 expression, in line with previous reports (Wang et al., 2002;Stein et al., 2004). In groups of slices treated with targeting Slc12a2 siRNA or Slc12a5 siRNA, we found a significant reduction of primary NKCC1 (n=9 slices; Mann-Whitney Rank Sum Test, U Statistic=10, P=0.014) or KCC2 (n=9 slices; Mann-Whitney Rank Sum Test, U Statistic=1.000, P<=0.001) antibody staining and protein expression relative to 4′,6-diamidino-2-phenylindole (DAPI) staining (Figure 8 c, d).

In all corresponding age-matched (DIV12-14) groups of organotypic hippocampal slices treated with siRNAs, electrical extracellular filed potential recording revealed spontaneous interictal- and ictal-like epileptiform discharges (Figure 8 e-j). In a control group of slices treated with non-targeting siRNA, bath application of WNK463 (1 µM for 30 min) progressively reduced the mean frequency of recurrent ILDs from 2.25±0.96 ILD/10 min in control ACSF (10 to 20 min) to 0.5±0.58 ILD/10 min (0 to 10 min over WNK463 application), and abolished recurrent ILDs (Figure 8e, f; n=4 slices; One Way RM ANOVA: DF=23, F(3, 15)=13.32, P<0.001; Tukey Test: Control vs WNK463 (0 to10 min over application), df=1.75, q=6.64, P=0.003; Control vs WNK463 (20 to 30 min over application), df=2.25, q=8.54, P<0.001). In slices treated with Slc12a2 siRNA (NKCC1), bath application of WNK463 (1 µM for 30 min) progressively reduced the mean frequency of recurrent ILDs from 1.75±0.5 ILD/10 min in control ACSF (10 to 20 min) to 0.25±0.5 ILD/10 min (10 to 20 min over WNK463 application), and abolished recurrent ILDs (Figure 8g, h; n=4 slices; One Way RM ANOVA: DF=23, F(3, 15)=6.29, P=0.002; Tukey Test: Control vs WNK463 (20 to 30 min over application), df=1.75, q=4.65, P=0.04). In slices treated with Slc12a5 siRNA (KCC2), bath application of WNK463 (1 µM for 30 min) progressively reduced the mean frequency of recurrent ILDs from 3±1 ILD/10 min in control ACSF (10 to 20 min) to 1±1 ILD/10 min during WNK463 application, and abolished recurrent ILDs (Figure 8i, j; n=3 slices; One Way RM ANOVA: DF=17, F(2, 10)=12.54, P<0.001; Tukey Test: Control vs WNK463 (0 to 10 min over application), df=2, q=5.1, P=0.04; Control vs WNK463 (20 to 30 min over application), df=3, q=7.61, P=0.004). Thus, siRNA targeted gene silencing of either Slc12a2 or Slc12a5 individually did not prevent or reduce the anti-ictal effects of WNK463, in line with pharmacological inhibition of NKCC1 and KCC2 (Figure 7).

### Simultaneous NKCC1 and KCC2 inhibition reduced the anti-ictal effects of WNK463

We next determined whether simultaneous inhibition of NKCC1 and KCC2 cotransport prevents the anti-ictal effects of WNK463 (Figure 9). Bath application of NKCC1 inhibitor bumetanide (10 µM) in combination with KCC2 inhibitor VU0463271 (1 µM) significantly reduced the mean frequency of ILDs, and consecutive application of WNK463 (1 µM) in the presence of combination of bumetanide and VU0463271 (Bum+VU) did not significantly reduce the mean frequency of ILDs (Figure 9b; n=5, One Way RM ANOVA: DF=74, F(4, 56)=6.1, P<0.001; Tukey Test: control compared to Bum+VU: df=2.8, q=5.07, P=0.04; control compared to WNK463 in the presence of Bum+VU: df=3.2, q=5.8, p=0.01; Bum+VU compared to WNK463 in the presence of Bum+VU: df=0.4, q=0.72, p=1.0). The corresponding power of electrical activity was reduced from a mean 955.25±335.72 µV^2^ in control to 885.16±169.25 µV^2^ during simultaneous bumetanide and VU0463271 application, and to 349.13±174.14 µV^2^ during consecutive application of WNK463 in the presence of bumetanide and VU0463271(Figure 9b; N=7, One Way RM ANOVA: DF=104, F(6, 84) =3.73, P<0.001; Tukey Test: control compared to Bum+VU, df=70.1, q=0.65, P=1.0; control compared to WNK463 in the presence of Bum+VU, df=606.12, q=5.6, P=0.012; Bum+VU compared to WNK463 in the presence of Bum+VU, df=536.03, q=4.95, P=0.04). Thus, simultaneous inhibition of NKCC1 and KCC2 did not prevent the anti-ictal effect of WNK463.

**Figure 9.**
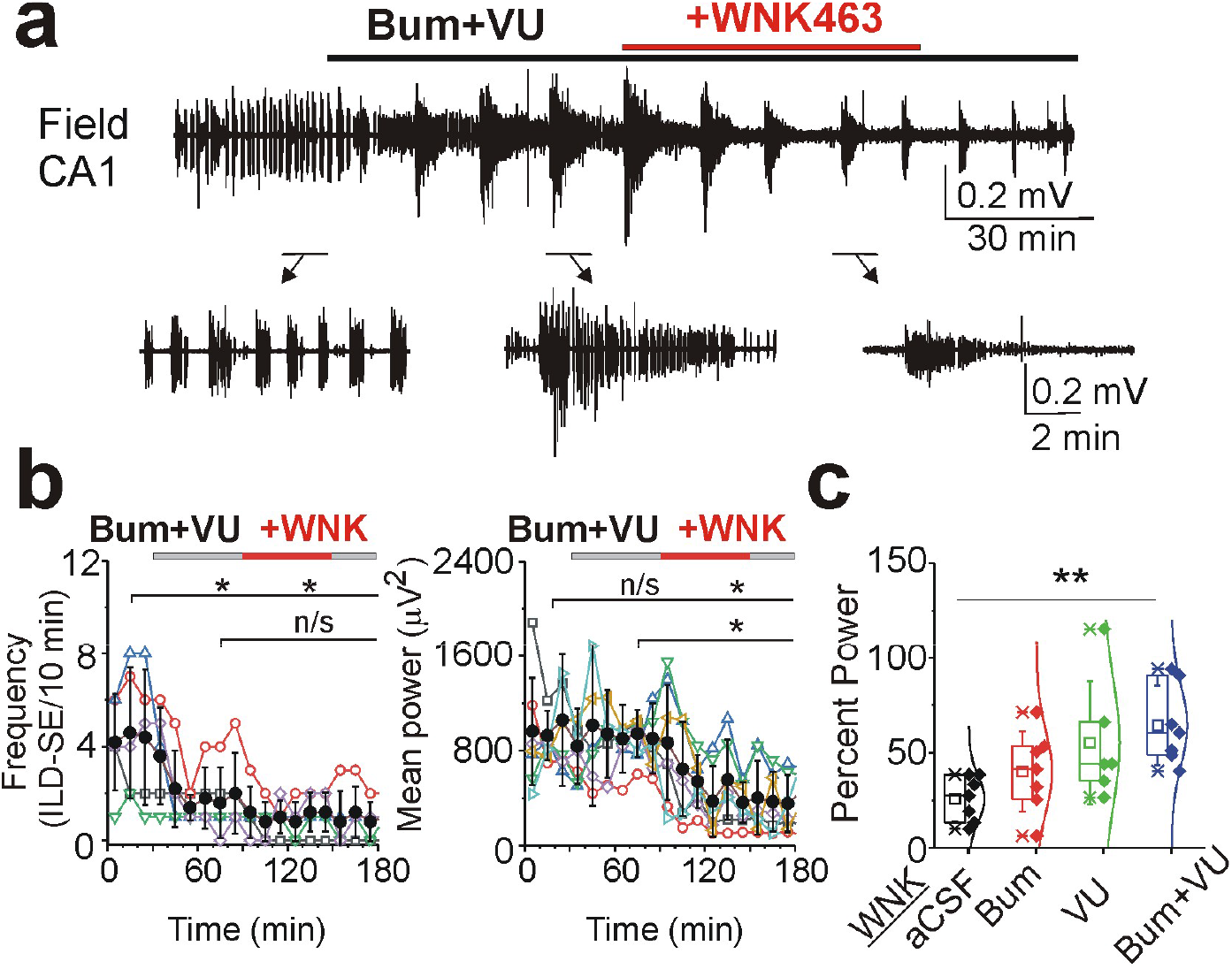
Simultaneous NKCC1 and KCC2 inhibition reduced the anti-ictal effects of WNK463. (**a**) Extracellular field potential recordings in the CA1 pyramidal cell layer in the organotypic hippocampal slices *in vitro*. WNK463 (1 µM) was applied in the presence of NKCC1 and KCC2 blockers bumetanide (10 µM) and VU0463271 (1 µM). Expansion of recurrent ILDs before and during drug applications. (**b**) WNK463 application in the presence of bumetanide in composition with VU0463271 did not change the frequency of ILDs but significantly reduced the mean power of epileptiform discharges preceding WNK463 application (*p<0.05; One-Way RM ANOVA, Tukey test). (**c**) Summary data of the effects of WNK463 on the mean power of electrical activity in control ACSF, in the presence of bumetanide (10 µM), in the presence of VU0463271 (1 µM), and in the composition of bumetanide (10 µM) and VU0463271 (1 µM). Simultaneous NKCC1 and KCC2 inhibition significantly reduced the anti-ictal effects of WNK463 (**P<0.01; One Way ANOVA, Holm-Sidak test).

We compared the mean effects of WNK463 on the power of electrical activity in control conditions with the corresponding effects of WNK463 in the presence of NKCC1 and KCC2 inhibitors (Figure 9c). On average, WNK463 significantly reduced the mean power of electrical activity to 25.77±11.8 percent of control, and the mean anti-ictal effects of WNK463 were not significantly different from the corresponding effects of WNK463 in the presence of NKCC1 inhibitor bumetanide (10 µM), or in the presence of KCC2 inhibitor VU0463271 (1 µM) (Figure 9c; One Way ANOVA, DF=26, F(3, 23)=4.05, P=0.019; Holm-Sidak method, WNK463+Bum compared to WNK463: df=14.3, t=1.21, P=0.24; WNK463+VU compared to WNK463: df=29.5, t=2.39, p=0.025). However, simultaneous inhibition of NKCC1 and KCC2 by the combination of bumetanide (10 µM) and VU0463271 (1 µM) significantly reduced the anti-ictal effects of WNK463 (Figure 9c; WNK463 in the presence of 10 µM Bumetanide+VU0463271 compared to WNK463: df=38.6, t=3.25, p=0.003). These results suggest that the anti-ictal effects of WNK-SPAK/OSR1 inhibitor WNK463 are mainly due to downstream interactions with NKCC1 inhibition and KCC2 activation, with additional effects due to interactions with an unidentified Cl^-^transporter, exchanger, or channel.

## Discussion

There is substantial interest in targeting the WNK-SPAK/OSR1-CCCs pathway for the therapy of various neurological disorders, including ischemic stroke (Josiah et al., 2021), neuropathic pain (Shekarabi et al., 2017), and anticonvulsant resistant seizures (Lee et al., 2022). We investigated the net effects of WNK463, an allosteric inhibitor of WNK-SPAK/OSR1 signaling, on [Cl^-^]_i_ elevation and extrusion rates during recurrent ILDs. We found that WNK463 improved [Cl^-^]_i_ homeostasis, progressively enhanced postictal [Cl^-^]_i_ extrusion rates, reduced ictal duration and abolished recurrent ILDs. Our investigations suggest that the powerful anti-ictal effects of WNK463 are primarily mediated by reduced NKCC1 and enhanced KCC2 transport activity that more efficiently prevent [Cl^-^]_i_ elevation and reduce activity-dependent increases in [Cl^-^]_i_ by suppression of ILDs.

Neuronal [Cl^-^]_i_ is a critical factor in determining whether postsynaptic GABA_A_-R mediated signaling is inhibitory or excitatory. In pathological conditions such as brain trauma, cytotoxic edema and/or recurrent seizures, in which [Cl^-^]_i_ may already be elevated, intense GABAergic input can further overwhelm the mechanisms that regulate chloride levels, leading to pathological increases in [Cl^-^]_i_ (Figures 1, 3-5). When [Cl^-^]_i_ increases, the E_GABA_ shifts towards more positive values, causing GABA-mediated depolarization and making neurons more likely to fire. This activity-dependent [Cl^-^]_i_ accumulation and a consequent shift from inhibitory to excitatory GABAergic signaling have been suggested to contribute to generation of ictal-like discharges (Fujiwara-Tsukamoto et al., 2006;Fujiwara-Tsukamoto et al., 2010;Ellender et al., 2014). In this fluorometric study, we did not measure RMP and E_GABA_ directly, but rather inferred the impact of electroneutral cation-chloride transport on these parameters from the effects on [Cl^-^]_i_ (Figure 2) and seizure activity (Figures 1, 3-9).

Restoration of [Cl^-^]_i_ equilibrium after synaptic activity is achieved by the net regulated activities of NKCC1 and KCC2 (Gamba, 2005). The NKCC1 membrane protein is the Na^+^-K^+^-2Cl^-^ transporter that under physiological conditions facilitates the uptake of Cl^-^ into cells, playing a crucial role in regulation of cell volume and [Cl^-^]_i_, especially during early brain development. NKCC1 facilitates seizures in the developing brain and pharmacological inhibition of NKCC1 enhances the efficacy of GABAergic anticonvulsants (Dzhala et al., 2005;Dzhala et al., 2008;Soul et al., 2021).Increased NKCC1 levels have also been observed in the subiculum of temporal lobe epilepsy patients (Palma et al., 2006;Huberfeld et al., 2007) and in several models of epilepsy, such as those induced by a reactive astrogliosis (Robel et al., 2015) or traumatic brain injury (Wang et al., 2017).

The KCC2 protein is a potassium-chloride co-transporter that maintains low [Cl^-^]_i_ (Rivera et al., 1999). KCC2 dysfunction has been implicated in epilepsy across human and experimental studies. Reduced KCC2 expression and impaired chloride extrusion have been demonstrated in resected tissue from patients with temporal lobe epilepsy, and pathogenic variants in SLC12A5 further support a causal role for impaired KCC2 function in human epilepsy (Huberfeld et al., 2007;Puskarjov et al., 2014). Experimental seizures can suppress KCC2 through several mechanisms in addition to WNK-SPAK/OSR1 signaling, including BDNF-TrkB-dependent downregulation, dephosphorylation of the stabilizing Ser940 site with subsequent internalization, and calpain-mediated proteolysis (Rivera et al., 2004;Silayeva et al., 2015;Puskarjov et al., 2012).These convergent mechanisms suggest that loss of KCC2 function may both result from and amplify seizure activity by weakening GABAergic inhibition. Accordingly, KCC2 has emerged as a therapeutic target: strategies include modulation of regulatory phosphorylation, enhancement of KCC2 expression, and direct pharmacological potentiation. Notably, the KCC2 activator OV350 enhances chloride extrusion and suppresses experimental seizures, including benzodiazepine-refractory status epilepticus, providing proof-of-concept for direct KCC2 activation as an antiseizure strategy (Jarvis et al., 2023).

Reduced KCC2 function or expression can contribute to activity-dependent accumulations of chloride and the excitatory shift in GABA signaling. Previous studies demonstrated the role of KCC2 transport activity in extrusion of [Cl^-^]_i_ and termination of ILDs (Sivakumaran et al., 2015;Dzhala and Staley, 2021). Our current data confirm that the high-affinity KCC2 inhibitor VU0463271 increased [Cl^-^]_i_ elevation and the duration of ILDs suggesting that inadequate KCC2 function contributes to the transition from the short ictal-like events to sustained *status epilepticus* (Figure 7). In contrast, activation of KCC2 transport activity efficiently recovers E_Cl_ during ILDs and restores GABAergic inhibition. These effects were correlated with more rapid termination of ILDs and electrical *status epilepticus* (Dzhala and Staley, 2021;Jarvis et al., 2023).

Hypoxia-ischemia, acute brain trauma, and recurrent seizures alter the equilibrium value of Cl^-^and impair the functional regulation of the CCCs, and are associated with cytotoxic cell swelling, acute and chronic accumulation of [Cl^-^]_i_, and GABA depolarizing responses, which foster seizures and anticonvulsant resistance via failure of inhibition (Dzhala et al., 2010;Dzhala et al., 2012). Antagonizing NKCC1 activity and/or stimulating KCC2 activity might reduce swelling and [Cl^-^]_i_ in injured neurons, restore GABAergic inhibition, suppress seizures, and prevent epileptogenesis.

NKCC1 and KCC2 are rapidly stimulated and inhibited, respectively, by direct phosphorylation mediated by the Cl^-^-sensitive WNK-SPAK/OSR1 complex (Alessi et al., 2014). The WNK family senses changes in the [Cl^-^]_i_, cell volume, extracellular osmolarity and transduces this information to CCCs (Shekarabi et al., 2017). WNKs stimulate the kinases SPAK and OSR1, which directly phosphorylate and stimulate NKCC1 or inhibit KCC2 (Kahle et al., 2013), providing compensatory and reciprocal regulation between transporters activity to maintain chloride homeostasis and neuronal excitability (Hartmann and Nothwang, 2022). Conversely, inhibiting the WNK-SPAK/OSR1 pathway might be an especially potent strategy to reduce neuronal Cl^-^ elevation and facilitate neuronal Cl^-^ extrusion by coincident NKCC1 inhibition and KCC2 activation. It was demonstrated that promoting KCC2 activity via WNK-SPAK/OSR1 inhibition can improve GABAergic inhibition and reduce neuronal excitability, potentially mitigating seizure activity (Lee et al., 2022).

Confusion may arise when discussing “baseline [Cl^-^]_i_” (Lee et al., 2022). In the presence of TTX, there is minimal synaptic GABA-mediated Cl^-^ loading, and the measured [Cl^-^]_i_ is the baseline [Cl^-^]_i_. During recurrent seizure activity, the [Cl^-^]_i_ measured between seizures is not likely to be at baseline due to ictal cytoplasmic Cl^-^ loading from GABA-mediated synaptic influx and incomplete cotransport of the excess [Cl^-^]_i_ out of the neuron. In this study, we found that the rate of Cl^-^transport out of the neuron was enhanced by blocking the WNK-SPAK/OSR1 pathway. Blocking this pathway did not change baseline [Cl^-^]_i_ significantly. However, blocking this pathway did reduce the interictal [Cl^-^]_i_ by enhancing the export of the ictal GABA-mediated Cl^-^ load. Considered from the point of Michaelis-Menten kinetics, changing the maximum velocity of Cl^-^cotransport does not alter the affinity of the transporter for [Cl^-^]_i_, which from this point of view would affect baseline [Cl^-^]_i_ (Staley and Proctor, 1999). Considered from the view baseline [Cl^-^]_i_ being set by Donnan exclusion (Glykys et al., 2014;Rahmati et al., 2021;Staley, 2024), altering the maximum velocity of Cl^-^ cotransport would also not be expected to alter baseline [Cl^-^]_i_.

## Limitations and future directions

Our experiments were conducted in organotypic hippocampal slices *in vitro*. Advantages of this preparation include homogenous injury and consistent epileptogenesis (Berdichevsky et al., 2012). This provides excellent control of the extracellular environment and imaging but does not replicate the *in vivo* vascular supply and maintenance of ionic homeostasis. While we find that increased WNK-SPAK/OSR1-mediated KCC2 activity via enhanced Cl^-^ extrusion rates robustly reduces seizure activity *in vitro*, it is possible that *in vivo*, increased KCC2 activity could lead to extracellular potassium accumulations (Viitanen et al., 2010) that would limit Cl^-^ extrusion independently of the velocity of KCC2 transport (Thompson et al., 1988;Staley and Proctor, 1999). Thus, it will be important to assess these findings *in vivo*.

## Materials and Methods

All animal-use protocols were in accordance with the guidelines of the National Institute of Health and the Massachusetts General Hospital Center for Comparative Medicine on the use of laboratory animals. All protocols were approved by the MGH Institutional Animal Care and Use Committee (IACUC).

### Culture of Organotypic Hippocampal Slices and Experimental Conditions

Transverse 350 μm hippocampal slices were prepared from C57BL/6, Clm1 (Duke University Medical Center, Durham, NC, USA) and sClm mice of either sex at postnatal (P) day 6-7 as previously described (Dyhrfjeld-Johnsen et al., 2010;Berdichevsky et al., 2012). Acute slices were mounted on poly-L-Lysine coated glass coverslips (Electron Microscopy Sciences, Hatfield, PA). Slices were incubated in 1000 μL of NeuroBasal/B27(1X) medium (Gibco by Life Technologies, Grant Island, NY) supplemented with 0.5 mM GlutaMAX and 30 μg/mL gentamicin (all from Invitrogen, Carlsbad, CA) in 6-well plates with low-evaporation lid (Becton Dickinson Labware, Franklin Lakes, NJ), in a humidified 37°C atmosphere that contained 5% CO_2_, placed on a rocking platform (< 1 cycle/min). Culture medium was changed bi-weekly. For acute recordings and imaging, slices were transferred to a submerged chamber and continuously superfused in oxygenated (95% O_2_ and 5% CO_2_) artificial cerebrospinal fluid (ACSF) containing (in mM): 126 NaCl, 3.5 KCl, 2 CaCl2, 1.3 MgCl_2_, 25 NaHCO_3_, 1.2 NaHPO_4_ and 11 Glucose (pH 7.4) at 32 ± 0.5°C and a flow rate of 2 mL/min. Slices were stabilized for 10-20 min in ACSF prior recordings and/or imaging. All organotypic hippocampal slices were used at Day *in Vitro* (DIV) 9-21. Slices from three to six (3–6) pups were used per each experimental group.

Pharmacological agents included bumetanide at a low concentration (10 μM) that blocks NKCC1, bumetanide at a high concentration (500 μM) that blocks NKCC1 and KCCs, the specific KCC2 blocker VU0463271 (1 μM), the GABA_A_ receptor (GABA_A_-R) antagonist SR95531 (10 μM), and the sodium voltage gated channel antagonist Tetrodotoxin (1 μM). Bumetanide and SR95531 were from Sigma-Aldrich CO, St. Louis, MO. VU0463271 and TTX were from TOCRIS Bioscience, Bristol, UK. WNK463 (1 μM) was from MedChemExpress, Monmouth Junction, NJ. Dimethyl sulfoxide (DMSO; 100 μl/100 ml) was used as an organic solvent (Dzhala and Staley, 2021).

### siRNA targeted genetic silencing of NKCC1 and KCC2

NKCC1 and KCC2 were selectively silenced using Dharmacom Accell™ (Dharmacon, Inc., Lafayette, CO) siRNA reagents in organotypic hippocampal brain slices at DIV 9-14. This siRNA technology does not require transfection reagents or instrument-based procedures that may result in loss of cell viability. Synthetic RNAi reagents were resuspended in RNAase-free solution (1x siRNA buffer diluted from 5x siRNA buffer (Item # B-002000-UB-100) using molecular grade RNAase-free water (Item # B-003000-WB-100)) according to siRNA resuspension protocol. Resuspended siRNA reagents were mixed with Accell delivery media (Item # B-005000-100) and used immediately. The final concentration was 1 µM siRNA per well in a 6-well plate. Culture medium was replaced with 1000 µL of the appropriate delivery mix (Accell siRNA and delivery media) to each well. Starting from DIV8-9, the organotypic slices were incubated in Accell siRNA delivery media at 37°C with 5% CO_2_ for 72 hours. As indicated by assay-dependent requirements (such as knockdown detection of a long-lived protein), Accell siRNA delivery media was changed back to culture medium and incubated at 37°C with 5% CO_2_ for an additional 24-72 hours following the standard 72 hours Accell incubation prior to assessing electrophysiological recordings and/or protein knockdown. Target sequence for ON-TARGET plus mouse Slc12a2 (NKCC1) siRNA (Item # L-044448-01-0020) was: (1) ACUAAGACAUAUCGACAGA, (2) CUAUGUAUGUUGUCGGAUU, (3) GAUUGUAAGAUCCGAGUAU, (4) AGGUCAAGCAAGACGUUAA; for mouse Slc12a5 (KCC2) siRNA (Item # L-058596-01-0020): (1) CGGCAUACACCUACGAGAA, (2) CAGGAGACAUCGCGUGUAU, (3) GAGCAAAGUUUCCGUUGAU, (4) UGGAGAUGCAUGAGAGCGA. Accell non-targeting siRNA (Item # D-001920-02-05) and Accell green (fluorescein) non-targeting siRNA (Item # D-001950-01-05) were used for control and siRNA delivery visualization.

Immunohistochemistry and confocal imaging were used for verification of siRNA targeted gene silencing (Bahari et al., 2024). Organotypic slices were prepared and processed for sequential immunostaining using a modified version of the protocol previously described in (Gogolla et al., 2006). The incubation time with primary antibody was extended to 3 nights, and 20% BSA in PBS was used as a blocking solution. The rabbit monoclonal anti-NKCC1 (1:200; AB303518-1007, lot # 1129040-4 from Abcam Inc, Waltham, MA) and mouse monoclonal anti-KCC2 (1:200; MABN88, lot # 4315985 from Millipore Sigma, Temecula, CA) were used as primary antibodies. The goat anti-rabbit Alexa Fluor 488 and goat anti-mouse Alexa Fluor 594 (1:1000; all from Invitrogen by ThermoFisher Scientific, Eugene, Oregon) were used as secondary antibodies.

Slices were washed and mounted onto glass microscope slides and covered by coverslip glass and mounting medium (Fluoromount-G, with DAPI, Invitrogen by ThermoFisher Scientific). Images were acquired on an Olympus FLUOVIEW FV3000 confocal laser scanning microscope and analyzed using ImageJ 1.53k software (Wayne Rasband and contributors, National Institutes of Health, USA). NKCC1 and KCC2 immunoreactivity (IR) were quantified within DG and CA1 regions of interest (ROIs). For each ROI, NKCC1 or KCC2 IR-positive area was normalized to the DAPI-positive area within the same region. Images were blinded, and a threshold for IR was determined across all images for each antibody. NKCC1-, KCC2-, and DAPI-positive areas were measured using the “Measure” function in ImageJ. Normalized IR was calculated as the NKCC1- or KCC2-positive area divided by the DAPI-positive area within the same DG or CA1 ROI.

### Electrophysiological recordings and data analysis

Extracellular field potentials were recorded in the CA3 and CA1 pyramidal cell layer of organotypic hippocampal slices using custom-made tungsten-coated 50 µm wire microelectrodes and ISO-DAM8A amplifier (WPI, Sarasota, FL). The electrical signals were digitized using an analog-to-digital converter DigiData 1322A (Axon Instruments, Inc, Union City, CA). AxoScope 10.7 and Clampfit 10.7 (Molecular Devices, San Jose, CA), Origin 2018 (OriginLab Corporation, Northampton, MA) and SigmaPlot 11.0 (Systat Software, Inc, San Jose, CA) programs were used for data acquisition and analyses. Recordings were sampled at 10 kHz. Interictal epileptiform discharges (IEDs) were defined as synchronous network-driven bursts characterized by short (0.1-3 s) duration and large amplitude population spikes. The frequency, duration, and amplitude of IEDs substantially varied between recurrent ictal-like discharges. Ictal-like discharges (ILDs) were defined as hyper-synchronous, large-amplitude and high-frequency population spikes followed by sustained ictal-tonic and/or intermittent ictal-clonic after-discharges, with the duration of the population spikes and after-discharge complex lasting more than 5 seconds, and subsequent post-ictal depression. *Status epilepticus* was defined as continuous ILDs for at least 5 min, or by sustained ictal-like tonic-clonic epileptiform discharges without recovery to baseline activity between the ILDs. Power spectrum analysis was performed on the electrical recordings after filtering with a Bessel high pass filter of 1Hz and applying a Hamming window function. The power of the electrical activity was calculated by integrating the root mean square value of the signal amplitude in corresponding time windows and frequency range from 1 to 1000 Hz. For comparison between slices, power was normalized for each slice with the highest value in control conditions. Experimental data were analyzed using a repeated measures ANOVA for two types of study design. Studies that investigated either (1) changes in mean (or median) parameters over three or more time points or consecutive time intervals); or (2) differences in mean (or median) parameters under three or more different conditions.

### Two-photon imaging of Clomeleon, quantitative and morphological analysis

Neuronal chloride concentration was determined in CA1 pyramidal neurons expressing the ratio-metric chloride indicators Clomeleon (Clm) (Kuner and Augustine, 2000) and superClomeleon (sClm) (Grimley et al., 2013;Rahmati et al., 2021). High-resolution two-photon excitation laser scanning imaging of the Cl^-^-sensitive yellow fluorescent protein (YFP) and the Cl^-^-insensitive cyan fluorescent protein (CFP) was performed on an Olympus Fluoview 1000 MPE microscope. A mode-locked titanium-sapphire laser (MaiTai, Spectra Physics) with 860 nm two-photon excitation was used to generate fluorescence. Emitted light passed through a dichroic mirror and was band-pass filtered through 480±15 nm (D480/30) for CFP and a 535±20 nm filter (D535/40) for YFP (FV10-MRCYR/XR). Time series acquisition of 720 frames (256×256 pixels for 254.46×254.46 µm) with 5-10 second intervals was performed to measure chloride concentration as a function of time in control conditions, during a 30-60 min period of applications of drugs, and over a 30-60 min period of wash-out. Transition from 10% above interictal [Cl^−^]_i_ to 90% of its maximal amplitude during onset of ILDs was used to estimate [Cl^−^]_i_ rise time. Mono-exponential fit (y=y0+A1*exp(–(x– x0)/t1) was used to estimate the decay time constant (t1) of [Cl^−^]_i_ recovery to baseline in individual cells.

For morphological analysis, organotypic slices were imaged through the CA1 pyramidal cell layer (Z-axis dimension 0-100 µm, 1-2 µm step size). ImageJ 1.51 software (National Institutes of Health) was used for quantitative analysis. Images were blinded and region of interests (ROIs) were selected using the chloride insensitive CFP fluorescence. The ratio of the YFP/CFP fluorescence intensity was used for [Cl^-^]_i_ calculation (Kuner and Augustine, 2000;Berglund et al., 2008;Glykys et al., 2009). The CFP emission of Clomeleon was used for the high-resolution morphological analysis (Dzhala et al., 2012).

### Statistical analysis

Group measures are expressed as mean ± standard deviation (SD) or median (25%-75%) as indicated. The Shapiro-Wilk test was used to determine normality of the data. The Student’s *t* test (paired or unpaired) was performed for parametric comparison of normally distributed data. The Wilcoxon Signed Rank Test (paired data), and Mann-Whitney test (unpaired data, two-tail) were used for non-parametric comparison of arbitrary distributed data. One-Way Repeated Measures Analysis of Variance (One Way RM ANOVA) was used for multiple comparison of parametric data to evaluate the differences in the mean values among the control and treatment groups. The Friedman RM ANOVA on Ranks was used for non-parametric data to determine the differences in the median values among the control and treatment groups. In a One Way RM ANOVA, (DF) represents the between-groups degree of freedom, and (F) represents the F-statistic and reported as F(DF_treatment_, DF_residual_)=value. In the Friedman RM ANOVA on Ranks, the chi-square statistic was used to assess the differences between the groups. The Tukey Test or Holm-Sidak Test was used for all pairwise comparisons of the responses to the different treatment groups. In the Tukey or Holm-Sidak test, (df) refers to the difference of means in the ANOVA, and (dr) refers to the difference of ranks in the ANOVA on Ranks. The level of significance was set at P<0.05.

## Acknowledgements

The authors acknowledge funding from the National Institutes of Health and National Institute of Neurological Disorders and Stroke grant R01NS120973 to VID and R35NS116852 to KJS.

## Funding Statement

The funders had no role in study design, data collection and interpretation, or the decision to submit the work for publication.

## Contributor Information

Volodymyr I Dzhala, Kevin J Staley

## Funding Information

This paper was supported by the following grants:

NIH and NINDS R01NS120973 to Volodymyr I Dzhala

NIH and NINDS R35NS116852 to Kevin J Staley

## Additional information

### Competing interests

No competing interests declared.

### Author contributions

DV - Conceptualization, Formal analysis, Validation, Investigation, Visualization, Methodology, Writing—original draft, Project administration, Writing—review and editing.

FHS, NR, AC, RB - Formal analysis, Validation, Investigation, Visualization, Methodology.

KK, KS - Conceptualization, Resources, Supervision, Funding acquisition, Project administration, Writing—review and editing.

### Ethics

Animal experimentation: All animal experimental procedures were performed in accordance with the National Institutes of Health Guide for the Care and Use of Laboratory Animals and approved by the MGH Institutional Animal Care and Use Committee (IACUC Protocol# 2020N000213).

### Data availability

All data generated or analyzed during this study are included in the manuscript.

## Notes

### Competing Interest Statement

The authors have declared no competing interest.

### Summary of Updates

This version of the manuscript has been revised to update the figure legend placement.

